# Host casein kinase 1-mediated phosphorylation modulates phase separation of a rhabdovirus phosphoprotein and virus infection

**DOI:** 10.1101/2021.11.16.468817

**Authors:** Xiao-Dong Fang, Qiang Gao, Ying Zang, Ji-Hui Qiao, Dong-Min Gao, Wen-Ya Xu, Ying Wang, Dawei Li, Xian-Bing Wang

**Affiliations:** State Key Laboratory of Agro-Biotechnology, College of Biological Sciences, China Agricultural University, Beijing 100193, China; College of Plant Protection, China Agricultural University, Beijing 100193, China

## Abstract

Liquid–liquid phase separation (LLPS) plays important roles in forming cellular membraneless organelles. However, how host factors regulate LLPS of viral proteins during negative-sense RNA (NSR) virus infections is largely unknown. Here, we used *Barley yellow striate mosaic virus* (BYSMV) as a model to demonstrate regulation of host casein kinase 1 in phase separation and infection of NSR viruses. We first found that the BYSMV phosphoprotein (P) formed spherical granules with liquid properties and recruited viral nucleotide (N) and polymerase (L) proteins *in vivo*. Moreover, the P-formed granules were tethered to the ER/actin network for trafficking and fusion. BYSMV P alone formed droplets and incorporated the N protein and genomic RNA *in vitro*. Interestingly, phase separation of BYSMV P was inhibited by host casein kinase 1 (CK1)-dependent phosphorylation of an intrinsically disordered P protein region. Genetic assays demonstrated that the unphosphorylated mutant of BYSMV P exhibited condensed phase, which promoted virus replication through concentrating the N, L proteins, and genome RNA into viroplasms. Whereas, the phosphorylation-mimic mutant existed in diffuse phase state leading to enhanced virus transcription. Collectively, our results demonstrate that host CK1 modulates phase separation of viral P protein and virus infection.

## Introduction

Over the last few years, increasing studies have shown that liquid–liquid phase separation (LLPS) has critical roles in assembly of cellular membraneless organelles such as P bodies, stress granules, cajal bodies, and the nucleolus (Handwerger *et al.*, 2005; Brangwynne *et al.*, 2009; Molliex *et al.*, 2015; Feric *et al.*, 2016; Boeynaems *et al.*, 2018; Darling *et al.*, 2019). LLPS concentrates specific molecules like proteins and nucleic acids into liquid-like compartments for fulfillment of their biological functions. The underlying molecular mechanisms have been of increased interests due to the important roles of LLPS in various physiological and pathological processes (Darling *et al.*, 2019). LLPS is usually triggered by intrinsically disordered regions (IDRs) of proteins and/or multivalent macromolecular interactions (Elbaum-Garfinkle *et al.*, 2015; Molliex *et al.*, 2015; Alberti *et al.*, 2018). In addition, LLPS is modulated by protein posttranslational modifications (PTMs), host factors, and cellular environment changes (Nott *et al.*, 2015; Banani *et al.*, 2017; Owen & Shewmaker, 2019).

Many negative-sense RNA (NSR) viruses are known to form membraneless replication compartments, called viroplasms, viral inclusion bodies (IBs), or viral factories (Lahaye *et al.*, 2009; Hoenen *et al.*, 2012; Rincheval *et al.*, 2017). Studies about animal NSR viruses have shown that LLPS plays important roles in viroplasm formation through concentrating viral and host components. The viroplasms of the rabies virus (RAPV) known as Negri bodies (NBs) were first reported to have the features of liquid organelles (Nikolic *et al.*, 2017). Subsequent studies have shown that another two animal NSR viruses, vesicular stomatitis virus and measles virus, exploit LLPS of viral proteins to form virus inclusion bodies (Heinrich *et al.*, 2018; Zhou *et al.*, 2019). The P protein of borna disease virus and the N protein of ebola virus are also sufficient to elicit formation of liquid organelles alone (Charlier *et al.*, 2013; Miyake *et al.*, 2020). However, most of these studies mainly focus on animal viruses (Brocca *et al.*, 2020; Su *et al.*, 2021), whereas it remains unclear in plant NSR viruses. More importantly, host factors regulating LLPS of viral viroplasms are largely unknown.

*Barley yellow striate mosaic virus* (BYSMV) is a member of the *Cytorhabdovirus* genus, family *Rhabdoviridae* in the order *Mononegavirales*. BYSMV infects cereal plants and severely affects crop production worldwide through persistent transmission by the small brown planthopper (SBPH, *Laodelphax striatellus*). The BYSMV genome encodes five structural proteins, including the nucleoprotein (N), phosphoprotein (P), matrix protein (M), glycoprotein (G), and polymerase (L), as well as another five accessory proteins, in the order 3′–N–P–P3–P4/P5–P6–M–G–P9–L–5′ (Yan *et al.*, 2015). Recently, we have developed minireplicon (BYSMV-agMR) systems and full-length cDNA clones of recombinant BYSMV (rBYSMV) for infections of plants and insects (Fang *et al.*, 2019; Gao *et al.*, 2019). Using the BYSMV reverse genetic systems, we have also shown that host factors, including casein kinase 1 and the deadenylation factor CCR4, are involved in virus cross-kingdom infections of host plants and insect vectors (Gao *et al.*, 2020; Zhang *et al.*, 2020). Interestingly, CK1 mediated phosphorylation of a highly serine-rich (SR) motif at the C-terminal intrinsically disordered region of the P protein regulates virus infection (Gao *et al.*, 2020). However, the mechanisms underlying regulation of the BYSMV P phosphorylation in virus infection are not well understood.

Here, using live-cell fluorescence microscopy to observe the localization of the BYSMV N, P and L core replication proteins, we noticed that ectopic expression of the BYSMV P protein alone resulted in spherical cytoplasmic granules. We also found that the BYSMV P-formed granules have properties of liquid organelles *in vivo* and *in vitro*. Host CK1-mediated phosphorylation of BYSMV P inhibits its LLPS. The roles of the BYSMV P phosphorylation and the LLPS in uncoupling virus replication and transcription were investigated and discussed.

## Results

### BYSMV P protein forms liquid-like granules through LLPS *in vivo*

To observe the BYSMV viroplasms *in vivo*, BYSMV-infected barley leaves were cut into ultra-thin sections and the structures formed in the cytoplasm were monitored by transmission electron microscopy. In agreement with animal negative-sense RNA (NSR) viruses (Hoenen et al., 2012; Rincheval et al., 2017), BYSMV infection induced formation of cytoplasm inclusions containing condensed ribonucleoproteins (RNPs) (Figure1A and Figure supplement 1). During rhabdovirus infection, the RNPs mainly consist of the N, P, and L proteins for replication and/or transcription (Jackson et al., 2005). To determine the core proteins involved in eliciting viroplasm formation, we examined the subcellular localization of ECFP-N, GFP-P, and L-mCherry in *N. benthamiana* leaves. At 2 days post infiltration (dpi), confocal imaging revealed that only GFP-P, rather than ECFP-N or L-mCherry, formed spherical granules throughout the cytoplasm (Figure 1B; Supplementary Video S1). Furthermore, ECFP-N and L-mCherry proteins were recruited into the GFP-P spherical granules, whereas CFP-N and L-mCherry failed to form condensates with free GFP (Figure supplement 2). These results suggest that BYSMV P is the scaffold factor for formation of spherical granules and then recruits the N and L proteins into condensed RNPs to facilitate viral infection.

**Figure 1.**
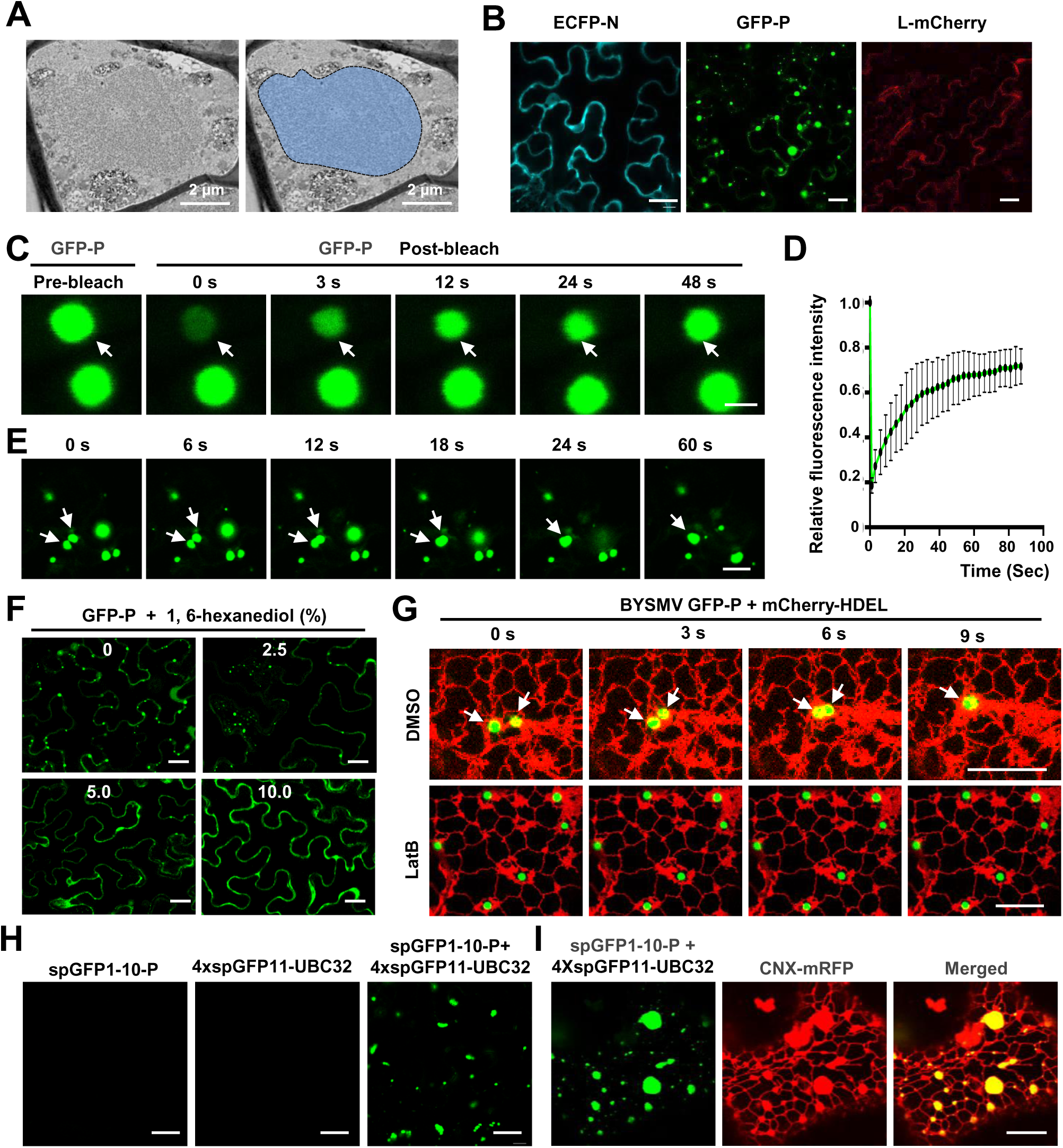
BYSMV P protein forms liquid-like granules through LLPS *in vivo*. **(A)** Transmission electron microscopy characterization of the BYSMV viroplasm in barley leaves infected by BYSMV at 10 dpi. The electron dense granular structure of the BYSMV viroplasm is highlighted by blue color in the right panel. Scale bars, 2 μm. **(B)** Confocal images showing subcellular distribution of ECFP-N, GFP-P, and L-mCherry in epidermal cells of *N. benthamiana* leaves at 2 dpi. Scale bars, 20 μm. **(C)** Representative FRAP images of GFP-P granules in epidermal cells of *N. benthamiana* leaves at 2 dpi. Leaves were treated with 10 μM LatB to inhibit movement of GFP-P granules at 3 h before photobleaching. Scale bar, 2 μm. **(D)** FRAP recovery curves of GFP-P granules. The intensity of each granule was normalized against their pre-bleach fluorescence. Data were presented as mean ± SD of 10 granules. **(E)** Confocal images showing fusion of two GFP-P granules in *N. benthamiana* leaf epidermal cells. White arrows indicate that GFP-P granules undergo fusion. Scale bar, 10 μm. **(F)** Representative images showing GFP-P localization after treatment with 0, 2.5%, 5.0%, 10.0% of 1, 6-hexanediol for 5 min in *N. benthamiana* leaf epidermal cells. Scale bars, 20 μm. **(G)** Time-lapse confocal micrographs showing the localization of GFP-P and mCherry-HDEL expressed in *N. benthamiana* leaf epidermal cells at 2 dpi. Leaves were treated with DMSO or 10 μM LatB at 3 h before imaging. White arrows indicate fusion of two GFP-P granules. Scale bars, 10 μm. **(H)** Confocal micrographs of *N. benthamiana* leaf epidermal cells expressed spGFP1-10-P, 4×SPGFP11-UBC32, or both at 2 dpi. Scale bars, 20 μm. (I) Confocal micrographs of *N. benthamiana* leaf epidermal cells expressed spGFP1-10-P, 4×SPGFP11-UBC32, and CNX-RFP at 2 dpi. The green signals indicate the contact sites of spGFP1-10-P granules with the tubular ER network. CNX-RFP is an ER marker. Scale bar, 10 μm.

We next used fluorescence recovery after photobleaching (FRAP) to determine whether the BYSMV GFP-P spherical granules have liquid properties. After photobleaching, approximately 80% of the granule GFP-P signals gradually recovered within 100 seconds (Figure 1C and d; Supplementary Video S2), indicating a rapid redistribution of the GFP-P protein between the membraneless granules and the surrounding cellular proteins. In addition, these GFP-P granules moved in the cytoplasm to fuse with each other (Figure 1E; Supplementary Video S1), and treatment with 1, 6-hexanediol, a chemical inhibitor of liquid-like droplets, efficiently dispersed the GFP-P granules (Figure 1F). Thus, these results demonstrate that the BYSMV P protein forms liquid-like granules through LLPS *in vivo*.

Emerging evidence shows that membrane-bound organelles provide platforms for assembly, fusion, and transport of membraneless granular condensates (Lee et al., 2020; Zhao and Zhang, 2020). To further evaluate the spatiotemporal dynamics of the GFP-P granules, mCherry-HDEL, a fluorescent ER marker, was monitored by fluorescence microscopy. Time-lapse confocal imaging analyses showed that the GFP-P granules are tethered tightly to the ER network and that their dynamics were correlated over time (Figure 1G; Supplementary Video S3). Treatment with the actin-depolymerizing agent latrunculin B (LatB) reduced both ER streaming and trafficking of the BYSMV-P granules (Figure 1G; Supplementary Video S4). Time-lapse confocal analyses also consistently showed that GFP-P granules moved along actin filaments marked by GFP-ABD2-GFP (Figure S3). These results suggest that the BYSMV-P granules move rapidly and fuse with each other during ER streaming in an actin-dependent manner.

Trafficking of GFP-P granules in close association with ER tubules strongly suggested that the GFP-P granules were tethered to ER tubules at molecular distances (10 to 30 nm) as membrane contact sites (MCSs) (Phillips and Voeltz, 2016). To examine the extent to which BYSMV P granules are tethered to the ER tubules, we used dimerization-dependent fluorescent protein domains to resolve the nanoscale resolution of GFP-P-ER contact in living cells. Previous studies have shown that two split GFP super-folder components (spGFP1-10 and spGFP11), can form functional spGFP green fluorescent signals when the two components interact at molecular distances (Pedelacq et al., 2006; Pedelacq and Cabantous, 2019). We fused a truncated form of the ER-localized ubiquitin conjugase UBC32 with 4×spGFP11 as an ER contact site marker (4×spGFP11-UBC32) (Cui et al., 2012; Li et al., 2020), and also fused the spGFP1-10 to the N terminus of P (spGFP1-10-P). At 2 dpi, neither 4×spGFP11-UBC32 nor spGFP1-10-P produced GFP signal when expressed alone (Figure 1H). Only co-expression of spGFP1-10-UBC32 and 4×spGFP11-P reconstituted a GFP signal that overlapped with the ER marker, CNX-RFP (Figure 1H and I) (Li et al., 2020), indicating that the GFP-P granules were tethered to the tubular ER network at molecular distances.

Collectively, our results suggest that the BYSMV P protein forms liquid spherical granules through LLPS *in vivo*. Furthermore, the ER/actin network provides a platform for dynamics of BYSMV-P granules. In agreement with the BYSMV GFP-P protein, the GFP-P protein of a closely related cytorhabdovirus, *Northern cereal mosaic virus*, had similar of liquid spherical granule features (Figure supplement 4; Supplementary Video S5 and S6).

### BYSMV P undergoes phase separation *in vitro*

To determine whether GFP-P liquid spherical granules are directly affected by the BYSMV P protein, we performed *in vitro* experiments to test phase separation of purified BYSMV P. To this end, we purified the recombinant proteins GFP and GFP-P from *E. coli* (Figure supplement 5A). As expected, the GFP-P protein, but not GFP, was able to condense into spherical droplets (Figure 2A; Figure supplement 5B). Moreover, treatment with 1, 6-hexanediol (HEX, 5%) efficiently dispersed the GFP-P droplets (Figure 2B). Moreover, increased GFP-P concentration and decreased NaCl concentration leads to enhancement of the numbers and sizes of the GFP-P droplets (Figure 2C).

**Figure 2.**
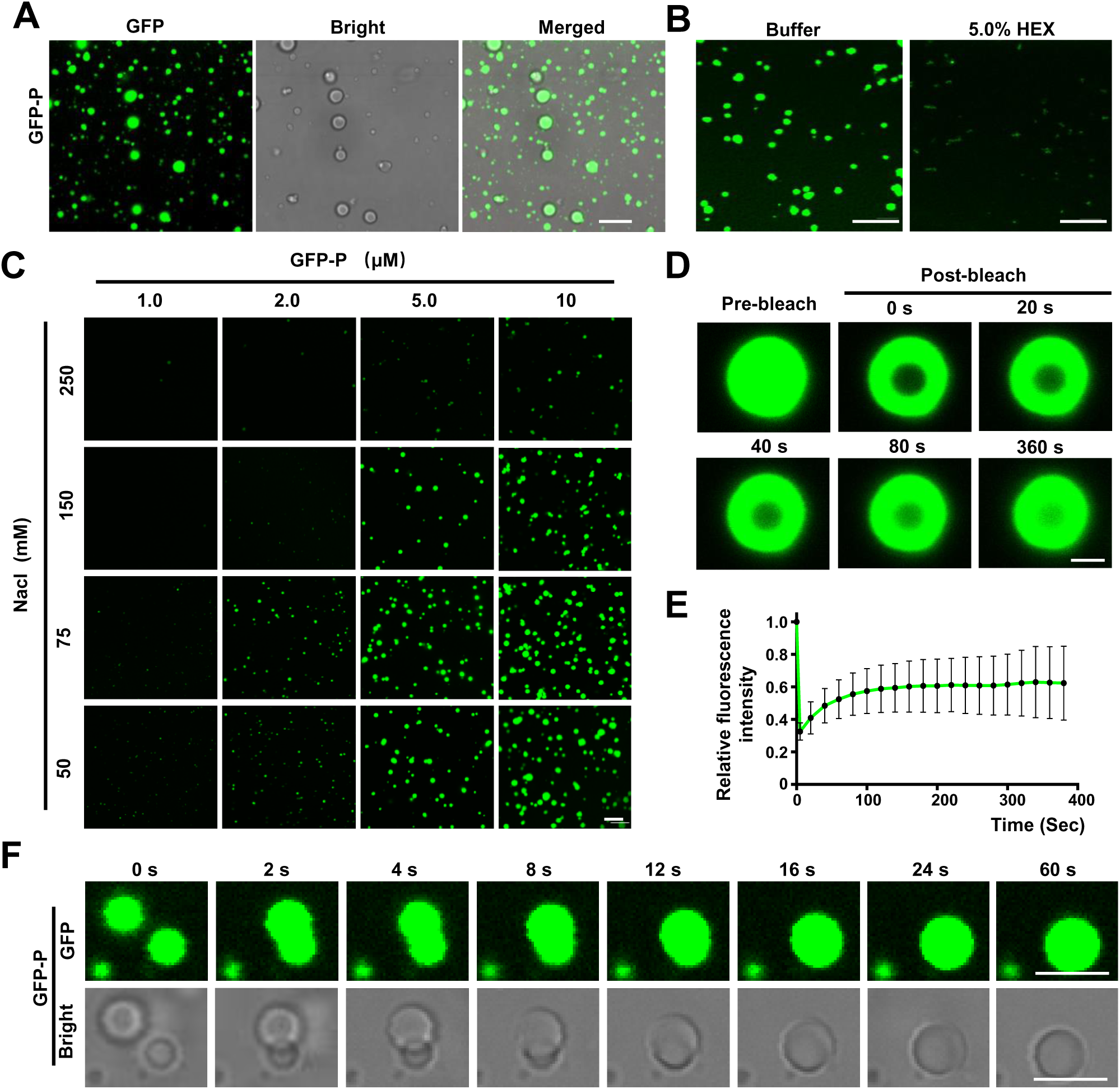
BYSMV P undergoes phase separation *in vitro*. **(A)** Confocal images showing that GFP-P formed droplets at the concentration of 10 μM in 125 mM NaCl. Scale bars, 10 μm. **(B)** Representative confocal images showing GFP-P droplets before or after treatment with 5.0% of 1, 6-hexanediol for 1 min. Scale bars, 10 μm. **(C)** Phase separation of GFP-P at different concentrations of GFP-P and NaCl. Scale bar, 20 μm. **(D)** Representative FRAP of GFP-P droplets *in vitro* at the concentrations of 10 μM in 125 mM NaCl. Scale bar, 1 μm. **(E)** FRAP recovery curve of GFP-P droplets. Data are shown as the mean ± SD of 10 droplets. **(F)** Representative images showing fusion of two GFP-P droplets *in vitro* at the concentration of 15 μM in 150 mM NaCl. Scale bars, 5 μm.

We also exploited FRAP to quantify the molecular dynamics within the GFP-P droplets by showing that ∼ 60% of the GFP-P signal in the droplets gradually recovered within 360 seconds after photobleaching (Figure 2D and E; Supplementary Video S7). We also observed that two approaching GFP-P droplets fused into a bigger droplet (Figure 2F; Supplementary Video S8). In summary, these *in vitro* results confirm that the BYSMV P protein alone has an intrinsic capacity for phase separation *in vitro*.

### P-formed droplets recruit the N protein and genomic RNA *in vitro*

*In vivo* assays showed that GFP-P granules were able to recruit the ECFP-N and L-mCherry proteins (Figure supplement 2). We next examined whether the BYSMV-P-formed droplets concentrated BYSMV N and genomic RNAs *in vitro*. The purified mCherry-N did not undergo LLPS *in vitro* (Figure 3A and Figure supplement 5B). However, when mCherry-N was incubated with GFP-P, the mCherry-N protein was gradually incorporated into GFP-P-formed droplets (Figure 3A). Interestingly, we observed three kinds of droplets *in vitro* (Figure 3B). Firstly, GFP-P and mCherry-N completely overlapped and were uniformly localized in the droplets (Figure 3B, top panel). Secondly, GFP-P is evenly distributed, but the mCherry-N signal decreased from the outside to the inside of the droplets (Figure 3B, middle panel). Thirdly, GFP-P and mCherry-N formed hollow droplets (Figure 3B, bottom panel).

**Figure 3.**
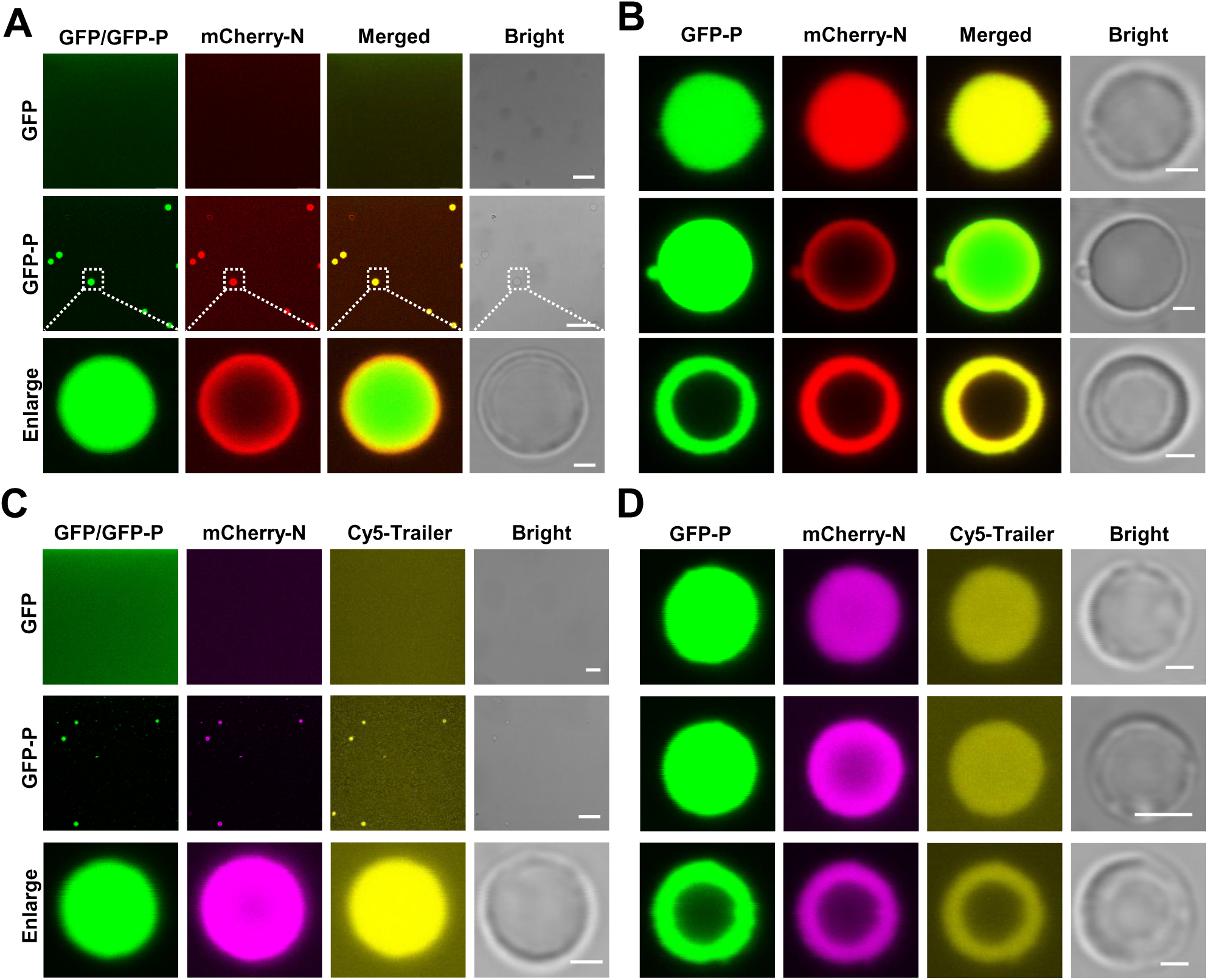
P-formed droplets recruit the N protein and genomic RNA *in vitro*. **(A)** Confocal images showing incorporation of mCherry-N into GFP-P droplets. Free GFP was unable to form droplets to recruit mCherry-N. Scale bars, 20 μm. Scale bars (enlarge panel), 1 μm. **(B)** Confocal images showing three representative droplet patterns formed by GFP-P and mCherry-N *in vitro*. Scale bars, 1 μm. **(C)** Confocal images showing incorporation of mCherry-N and Cy5-trailer into GFP-P droplets. In contrast, free GFP was unable to form droplets or recruit mCherry-N and Cy5-labeled trailer. Scale bars, 20 μm. Scale bar (enlarge panel), 1 μm. **(D)** Confocal images showing three representative droplet patterns formed by GFP-P, mCherry-N, and Cy5-trailer *in vitro*. Scale bars, 1 μm.

To test whether the GFP-P and mCherry-N droplets can recruit genomic RNA, a 334 nt RNA fragment corresponding to the 5′ trailer of BYSMV negative RNA genome was labeled by Cy5 (Cy5-Trailer). The Cy5-Trailer fragment was added to GFP-P/mCherry-N or GFP/mCherry-N mixtures *in vitro*. As expected, both mCherry-N and Cy5-Trailer were incorporated into the GFP-P droplets, whereas they appeared to be evenly distributed when incubated with GFP (Figure 3C). Moreover, the GFP-P/mCherry-N/Cy5-Trailer mixture formed three distinct kinds of droplets including uniform, hollow, and intermediate condensates (Figure 3D). Taken together, these results suggest that BYSMV P-formed liquid droplets can incorporate BYSMV N and genomic RNA *in vitro*.

### Phase separation of BYSMV P is inhibited by P protein phosphorylation

BYSMV P is a phosphoprotein whose phosphorylation states affect virus replication and transcription (Gao et al., 2020). *In silico* predictions via PONDR suggest that the BYSMV P protein contains three intrinsically disordered regions (IDRs) (Figure 4A). Interestingly, five highly phosphorylated Ser residues (amino acids 189, 191, 194, 195, and 198) are present in an Serine-rich (SR) motif (^189^SASRPSSIAS^198^) located in the middle IDR of BYSMV P (Gao et al., 2020). Given that protein IDRs are usually involved in phase separation (Brangwynne et al., 2015), we hypothesized that the phosphorylation states of the BYSMV SR region might affect the phase separation ability of BYSMV P. To test this hypothesis, we carried out site-directed mutagenesis to replace each Ser residue in the middle IDR (IDR2) with an Ala (GFP-P^S5A^) or Asp residues (GFP-P^S5D^) to mimic unphosphorylated and hyperphosphorylated states, respectively (Figure 4A). The GFP-P^WT^, GFP-P^S5A^, and GFP-P^S5D^ proteins were individually expressed in *N. benthamiana* leaves by agroinfiltration. Intriguingly, in contrast to GFP-P^WT^ (∼116 granules per view), GFP-P^S5A^ formed relative fewer (∼70 granules per view) but larger granules, whereas GFP-P^S5D^ formed very fewer granules (∼18 granules per view) and more evenly located in the cytoplasm (Figure 4B and C). Statistical analyses revealed that the sizes of less than one-fifth of the GFP-P^WT^ granules were larger than 1 μm in diameter, whereas approximately one-third GFP-P^S5A^ granules had larger diameters (Figure 4D).

**Figure 4.**
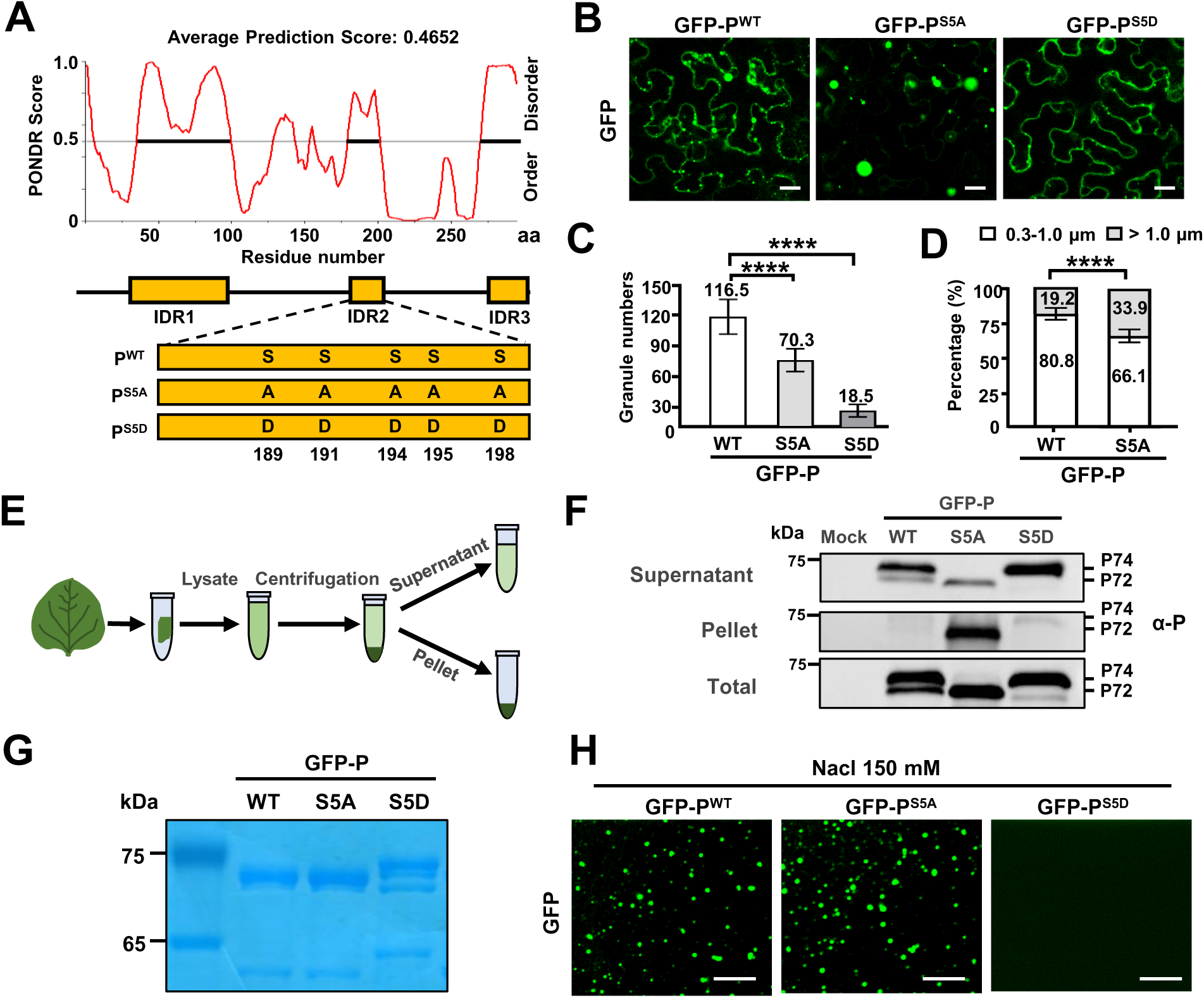
Phase separation of BYSMV P is inhibited by P protein phosphorylation. **(A)** The predicted IDRs of BYSMV P and schematic representation of its phosphorylation mutants (P^S5A^ and P^S5D^). IDRs were predicted according to the online tool PONDR and indicated by yellow boxes. **(B)** Confocal images showing subcellular distribution of GFP-P^WT^, GFP-P^S5A^, and P-GFP^S5D^ in *N. benthamiana* leaf epidermal cells at 2 dpi. Scale bars, 20 μm. **(C)** Statistical analyses of GFP granules from eight epidermal cells of *N. benthamiana* leaves expressing GFP-P^WT^, GFP-P^S5A^, or GFP-P^S5D^. **(D)** Statistical diameter analyses of GFP-P^WT^, GFP-P^S5A^ granules. **(E)** Work flow showing granule sedimentation assays using *N. benthamiana* leaves expressing GFP-P^WT^, GFP-P^S5A^, or P-GFP^S5D^ at 2 dpi. **(F)** Western blotting analyses of supernatant, pellet, and total proteins isolated in panel E. **(G)** SDS-PAGE showing purified GFP-P^WT^, GFP-P^S5A^, or P-GFP^S5D^ purified from *E. coli*. **(H)** Confocal images showing droplet formed by GFP-P^WT^, GFP-P^S5A^, or P-GFP^S5D^ *in vitro*. Scale bar, 10 μm. In C and D, all results were obtained from eight views (175 μm × 175 μm). Error bars indicate SD. **** *p* < 0.0001 (Student’s t-test).

In previous studies, P-bodies can be isolated from animal cells by centrifugation and fluorescence activated particle sorting (Hubstenberger et al., 2017). To further examine condensed or diffuse states of GFP-P^WT^, GFP-P^S5A^, and GFP-P^S5D^, total extracted protein samples of infiltrated leaves were centrifuged at 10,000 × g for 10 min, and the supernatant and pellet fractions were subjected to Western blotting analyses with anti-P antibodies (Figure 4E). Interestingly, two bands of GFP-P^WT^ corresponding to about 72 kD (P72) and 74 kD (P74) were present, whereas GFP-P^S5A^ and GFP-P^S5D^ existed as P72 and P74, respectively (Figure 4F). As expected, the GFP-P^S5D^ protein was mainly concentrated in the supernatant fraction, while the GFP-P^S5A^ protein was mainly in the pellet fraction (Figure 4F), indicating that GFP-P^S5D^ and GFP-P^S5A^ primarily existed as soluble and condensed states, respectively.

We further examined *in vitro* phase separation of the purified recombinant GFP-P^WT^, GFP-P^S5A^, and GFP-P^S5D^ proteins (Figure 4G). The results were consistent with the *in vivo* results (Figure 4B), as GFP-P^WT^ and GFP-P^S5A^, but not GFP-P^S5D^, underwent phase separation in 150 mM NaCl (Figure 4H). Assays at different protein concentrations in 125 mM NaCl indicated that the dephosphorylation state of GFP-P^S5A^ had enhanced phase separation activity compared with GFP-P^WT^ (Figure supplement 6A). Taken together, these results confirm that phosphorylation of the SR region within the middle IDR of BYSMV P significantly impairs phase separation *in vivo* and *in vitro*.

### Host CK1 negatively regulates BYSMV P phase separation

Given the conserved CK1 kinases among host plants and insect vectors that directly target the five Ser residues of the SR motif *in vivo* and *in vitro* (Gao et al., 2020), it would be interesting to determine whether CK1 affects BYSMV P phase separation *in vivo*. To this end, GFP-P^WT^, GFP-P^S5A^, and GFP-P^S5D^ were expressed with the empty vector (EV) or the CK1 proteins (NbCK1) in *N. benthamiana* leaves. As expected, the GFP-P^WT^ granules were almost dispersed upon co-expression of NbCK1 (Figure 5A). By comparison, GFP-P^S5A^ granules were not affected by NbCK1 (Figure 5A). Again, GFP-P^S5D^ was defective in granule formation in either the presence or absence of NbCK1.3 (Figure 5A). Western blotting analyses showed that co-expression of NbCK1 drastically increased the hyper-phosphorylated P74 form of GFP-P^WT^ compared with equal accumulation of P72 and P74 forms during co-expression of GFP-P^WT^ and EV (Figure 5B, compare lane 2 and 3). In contrast, the P72 form of GFP-P^S5A^ and the P74 form of GFP-P^S5D^ were not affected by co-expression of either NbCK1 or EV (Figure 5B). These results indicate that host NbCK1 mainly inhibits BYSMV P phase separation by phosphorylating the five Ser residues of the SR motif within the middle IDR region of BYSMV P (Figure 4A).

**Figure 5.**
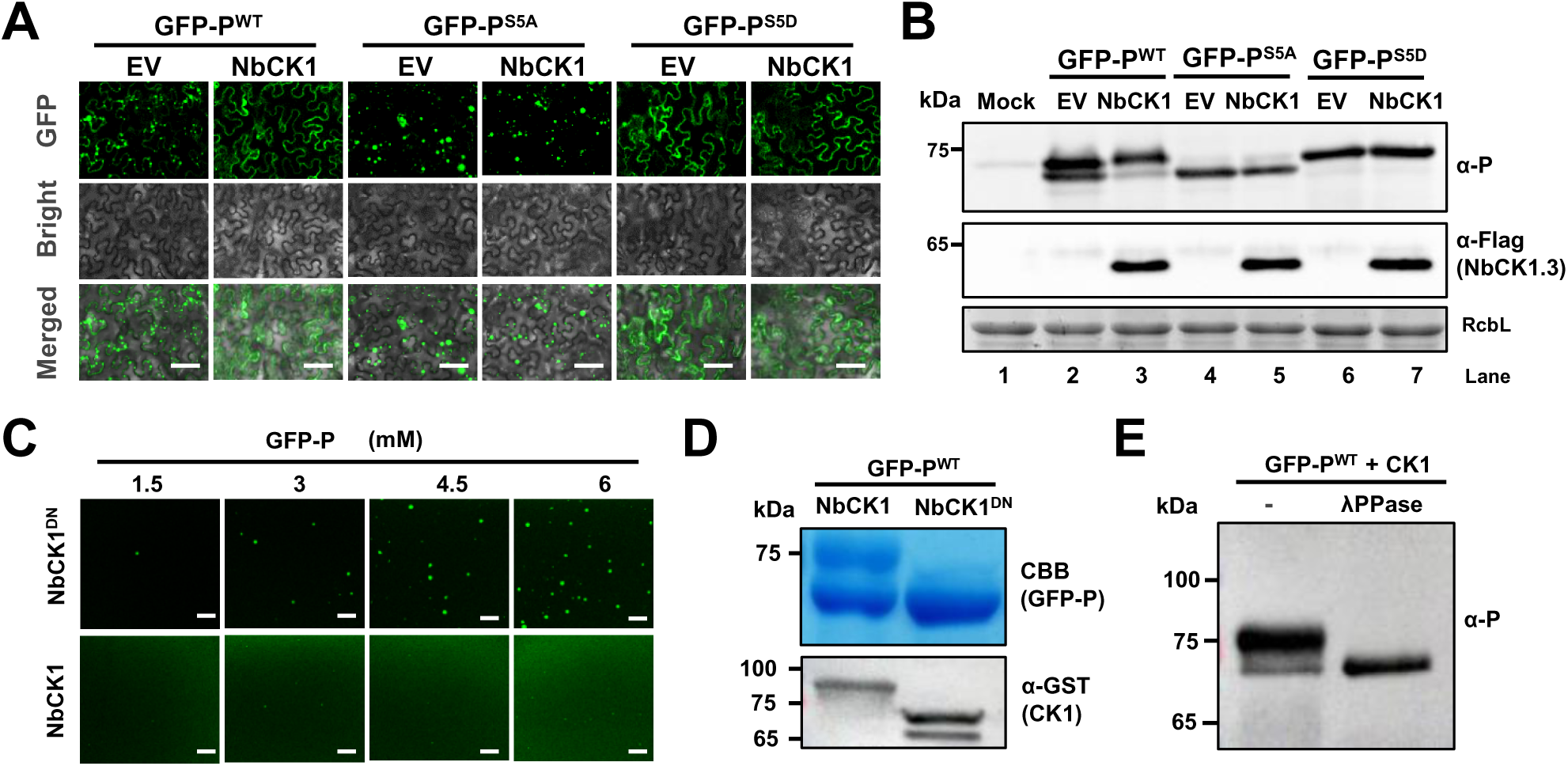
Host CK1 inhibits phase separation of BYSMV P *in vivo* and *in vitro*. **(A)** Confocal images showing subcellular distribution of GFP-P^WT^, GFP-P^S5A^, and P-GFP^S5D^ co-expressed with empty vector (EV) or NbCK1 in *N. benthamiana* leaf epidermal cells at 2 dpi. Scale bars, 50 μm. **(B)** Western blotting detecting accumulation of GFP-P^WT^, GFP-P^S5A^, and P-GFP^S5D^ in the leaves as shown in panel A. **(C)** Confocal images showing droplet formation of GFP-P^WT^ purified from *E. coli* co-expressing NbCK1 or NbCK1^DN^. GFP-P^WT^ was diluted to different concentration and 125 mM NaCl. Scale bars, 20 μm. **(D)** SDS-PAGE showing purified GFP-P^WT^, GFP-P^S5A^, or P-GFP^S5D^ in the samples of panel C. Expression of GST-tagged NbCK1 or NbCK1^DN^ was examined by Western blotting analyses with anti-GST antibodies. **(E)** Western blot detecting GFP-P^WT^ treated with lambda protein phosphatase (λ-PPase) or mock buffer (-) with anti-P antibodies.

Since eukaryotic-type protein kinases are absent in *E. coli*, we used a bacterial co-expression system to isolate phosphorylated BYSMV P protein from *E. coli*. We then co-expressed GFP-P^WT^ with NbCK1 or its loss of function mutant (K38R and D128N, NbCK1^DN^) (Gao et al., 2020). The purified GFP-P^WT^ protein underwent phase separation when co-expressed with NbCK1^DN^, but co-expression of NbCK1 severely inhibited phase separation of GFP-P^WT^ (Figure 5C). In addition, the SDS-PAGE gel showed that co-expression of NbCK1, rather than NbCK1^DN^, resulted in production of the upper P74 band of GFP-P^WT^ (Figure 5D). Moreover, we failed to detect the P74 band of GFP-P^WT^ after λ-protein phosphatase (λPPase) treatment *in vitro* (Figure 5E), indicating that P74 represents hyper-phosphorylated forms of GFP-P^WT^ elicited by co-expressed NbCK1. In agreement with NbCK1, the CK1 orthologue of barley plants (HvCK1) suppressed phase separation of GFP-P^WT^ in co-expression assays (Figure supplement 6B and S6C).

Collectively, host CK1 proteins inhibit phase separation of BYSMV P by phosphorylating the SR region of the BYSMV P middle IDR.

### Condensed phase of BYSMV P facilitates virus replication but inhibits transcription

We next investigated the relevance of BYSMV P phase separation on replication and transcription. Recently, we have developed a BYSMV minireplicon (BYS-agMR) to mimic viral replication and transcription processes (Fang et al., 2019). Based on the pBYS-agMR plasmid, we generated a frame-shift vector (pBYS-agMR-RFP) to abolish translation of the GFP mRNA (Figure supplement 7A). Then, we used GFP-P^WT^, GFP-P^S5A^, or GFP-P^S5D^ to rescue BYS-agMR-RFP in *N. benthamiana* leaves after co-agroinfiltration of pBYS-agMR-RFP, pGD-VSRs, pGD-N, and pGD-L (Figure supplement 7). Consistent with the result above (Figure 4B), GFP-P^S5A^ formed larger spherical granules than those of GFP-P^WT^, whereas GFP-P^S5D^ was diffuse in the cytoplasm (Figure 6A). Furthermore, the numbers of RFP foci in the GFP-P^S5A^ samples were significantly higher than those of GFP-P^WT^, whereas GFP-P^S5D^ expression resulted in a reduced number of RFP foci (Figure 6A). Western blotting analyses consistently showed that RFP accumulation was highest after GFP-P^S5A^ expression, followed by GFP-P^WT^ and GFP-P^S5D^ (Figure 6B).

**Figure 6.**
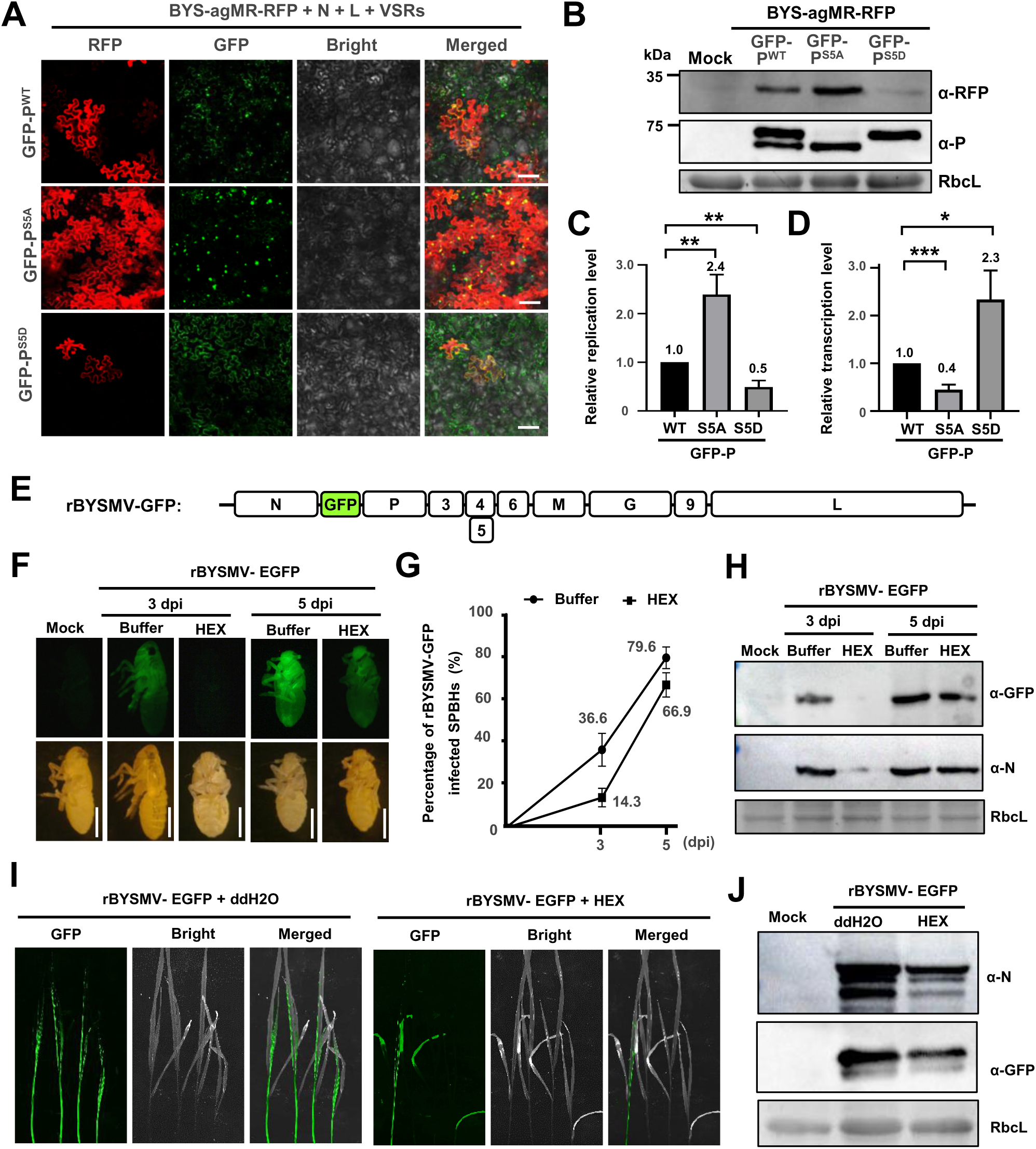
Phase separation of GFP-P modulates virus replication and transcription. **(A)** RFP foci in *N. benthamiana* leaves infiltrated with *Agrobacterium* for co-expression of BYS-agMR-RFP, N, L, and VSRs with GFP-P^WT^, GFP-P^S5A^, or GFP-P^S5D^ at 8 dpi. Scale bars, 100 μm. **(B)** Western blotting analyzing accumulation of RFP, GFP-P^WT^, GFP-P^S5A^, and GFP-P^S5D^ in the leaf samples of panel A. **(C)** Quantitative real-time PCR analyzing the relative levels of minigenome replication supported by the GFP-P^WT^, GFP-P^S5A^, and GFP-P^S5D^ proteins. **(D)** Quantitative real-time PCR analyzing the relative levels of RFP mRNA in the same samples of A. **(E)** Schematic diagram of the rBYSMV-GFP genome. A *gfp* gene was inserted between N and P genes to express GFP reporter and indicate virus systemic infection. **(F)** The images of SBPHs infected by rBYSMV-GFP with 3% HEX or ddH2O. At 3- and 5- dpi, the insects were observed with fluorescence stereo microscope. Scale bars, 1 mm. **(G)** Effects of 3% HEX or buffer on the infection rates of rBYSMV-GFP in SBPHs at 3- and 5- dpi. n > 35. **(H)** Western blot analyses of expression of GFP and BYSMV N proteins in SBPHs shown in F with specific antibodies against GFP and BYSMV N proteins, respectively. **(I)** Systemic infection of rBYSMV-GFP in barley plants treated with buffer or 3% HEX at 10 dpi. Scale bars, 2 cm. **(J)** Western blotting analyzing accumulation of BYMV N and GFP in the barley plants as shown in panel I. In C and D, all results were obtained from three independent biological repeats. The values in the GFP-P^WT^ samples were set to 1. The *EF-1A* gene was used as a negative control. Error bars indicate SD. * *p* < 0.05, ** *p* < 0.01, and *** *p* < 0.001 (Student’s t-test).

We subsequently performed RT-qPCR to compare relative accumulation of RNA products of virus replication and transcription. Accumulation of the full-length genome minigenome was drastically upregulated in GFP-P^S5A^ samples but decreased in GFP-P^S5D^ samples compared with those of GFP-P^WT^ samples (Figure 6C). Considering that the replicating products are the templates for virus transcription, the relative transcription activities were measured based on the ratios of mRNA accumulation to full-length minigenomic RNA. Compared with GFP-P^WT^, GFP-P^S5D^ supported higher transcription activity but GFP-P^S5A^ expression significantly compromised transcription (Figure 6D). Collectively, GFP-P^S5A^ with increased LLPS activity supports enhanced virus replication but decreased transcription. In contrast, GFP-P^S5D^ with severely impaired LLPS activity inhibited virus replication but facilitated transcription.

BYSMV can cross-kingdomly infect host plants and insect vectors. We next examined effect of BYSMV P phase separation on virus infection in its insect vector, the small brown planthopper (SBPH; *Laodelphax striatellus*). To this end, we coinjected the thoraxes of SBPHs with crude extracts of rBYSMV-GFP (Figure 6E) and 1, 6-hexanediol (3%) to inhibit phase separation. At 3- and 5- dpi, 1, 6-hexanediol (HEX)-treated insects exhibited lower GFP fluorescence signal (Figure 6F) and reduced infection ratios (Figure 6G) than those treated with buffer. In consistence, immunoblot analyses revealed that accumulation of GFP and N proteins was lower in HEX-treated insects than those treated with buffer (Figure 6H). We also treated rBYSMV-GFP-infected barley leaves with 3% HEX to inhibit phase separation of BYSMV P during virus infection. At 10 dpi, GFP fluorescence signal derived from rBYSMV-GFP was obviously less intense in HEX-treated barley plants than those of mock treatment (Figure 6I). Accumulation of GFP and N proteins was consistently reduced after HEX treatment (Figure 6J). Collectively, these results indicate that phase separation is important for cross-kingdom infection of BYSMV in insect vectors and host plants.

In summary, the unphosphorylated BYSMV-P protein undergoes LLPS and forms spherical granules as scaffold centers that concentrate the N and L proteins and genomic RNA to promote virus replication. In contrast, the conserved CK1 protein kinases phosphorylate the SR region of BYSMV P to hyper-phosphorylated P forms and compromise LLPS, which results in soluble RNPs that promote viral transcription (Figure 7). Therefore, the BYSMV P phosphorylation states modulate phase separation and uncouple replication and transcription, which plays important roles in virus cross-kingdom infection in host plants and insect vectors.

**Figure 7.**
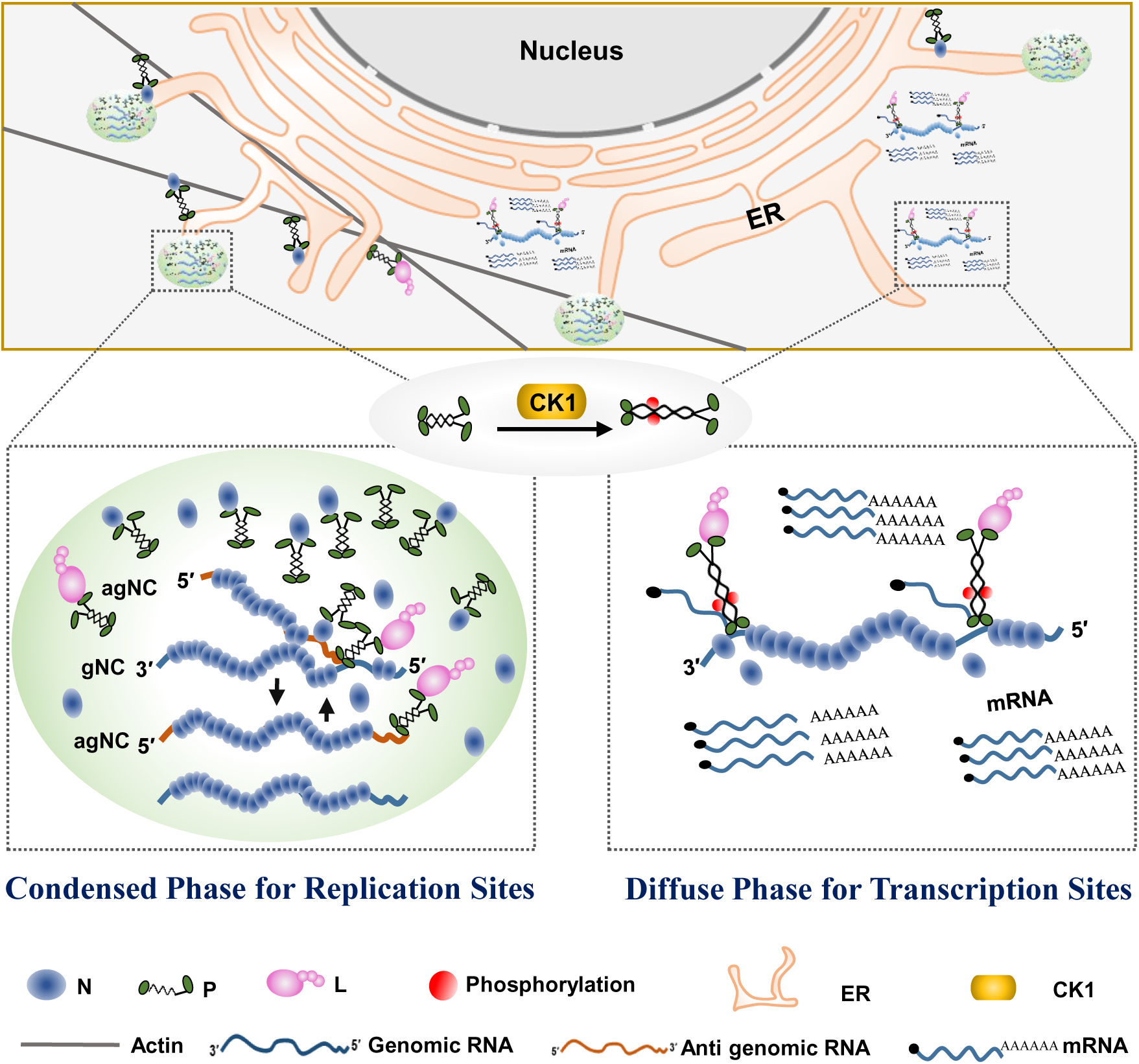
Model for phase separation of BYSMV P in uncoupling rhabdovirus replication and transcription. Rhabdovirus replication requires high concentration of viral N protein for encapsidating newly synthesized genomic/antigenomic RNA. Thus, unphosphorylated BYSMV P undergoes phase separation, and then recruits the N and L proteins, as well as genomic RNAs into membraneless condensates for optimal replication. In addition, the granules move along the ER/actin network and fuse with each other. In the transcription sites, the SR motif in the middle IDR of BYSMV P is phosphorylated by CK1, and the resulting hyper-phosphorylated P is unable to undergo phase separation, which facilitates virus transcription and viral mRNA release for viral protein translation.

## Discussion

Membraneless virus inclusion bodies or viroplasms are hallmarks of NSR virus infections. In recent studies, LLPS has emerged as a critical mechanism in formation of replication factories of animal NSR viruses (Brocca *et al.*, 2020; Su *et al.*, 2021). In contrast, the mechanisms whereby plant NSR viruses form viroplasms as replication sites have not been characterized. Here, we provide the first evidence, to the best of our knowledge, that a plant NSR virus uses LLPS to form viral replication proteins and genomic RNA condensates. We have identified a novel cytorhabdovirus P protein function that provides a scaffold protein for viroplasms formation. The BYSMV P protein can undergo phase separation alone *in vivo* and *in vitro* to concentrate the N and L proteins, and the viral RNA into membraneless granules that promote virus replication.

Another intriguing finding in our study is that host CK1-dependent phosphorylation of the BYSMV P protein inhibits phase separation. Our previous study has shown that conserved CK1 kinases of host plants and insect vectors are responsible for phosphorylation of a serine-rich (SR) motif at the middle IDR region of BYSMV P (Gao *et al.*, 2020). In the present study, we found that the BYSMV P condensates were dispersed by phosphorylation mimic mutations (P^S5D^) of the SR motif or overexpression of CK1 proteins *in vivo* (Figure 4B and 5A). These results are in agreement with recent reports in mammalian cells in which casein kinase-2 (CK2)-mediated phosphorylation of ZO (zona occludens), a cytoplasmic scaffolding protein, inhibits homologous phase separation (Beutel *et al.*, 2019). The partitioning defective 3/6 (Par3/Par6) condensates are also dispersed by PKC-mediated phosphorylation (Liu *et al.*, 2020). Thus, precisely balanced phosphorylation states may represent additional regulators of protein phase separation and cellular functions.

Rhabdovirus transcription and replication complexes contain common virus replication derivatives, in which the N-RNA complex serves as a template for viral RdRp complexes comprising the P and L proteins (Ivanov *et al.*, 2011). Therefore, models to explain uncoupling of rhabdovirus transcription and replication processes have remained elusive for many years because of the similarity of the N, P and L proteins and the viral genomic RNA components. Here, we demonstrate that the phosphorylation mutants (P^S5A^) in the SR motif of BYSMV P have high LLPS activity that is coupled with enhanced virus replication. On the other hand, the phosphorylation-mimic mutant (P^S5D^) is impaired in LLPS, which facilitates virus transcription but significantly inhibits virus replication. Therefore, these results suggest that condensed phase of RNPs as viroplasms are a positive regulator for virus replication, but inhibit virus transcription. During rhabdovirus replication, the newly synthesized genome or antigenome is encapsidated by the N protein, which requires a continuous supply of RNA-free N molecules. Therefore, it is not surprising to find that LLPS of BYSMV P facilitates concentration of the BYSMV N protein into virus replication sites. In contrast, viral mRNAs are released without N encapsidation, which results in dispersed protein rather than condensed N protein in virus transcription sites (Figure 7). Therefore, we propose that the phosphorylation of the BYSMV P SR region by host CK1s abrogates the liquid-like membraneless viroplasms and subsequently drives the switch from virus replication to transcription.

Emerging evidence shows that the biogenesis and dynamics of phase-separated membraneless condensates require membranes as assembly platforms for transportation (Zhao & Zhang, 2020). For instance, ER contact sites regulate the biogenesis and fission of two important membraneless organelles, processing bodies (PBs) and liquid spherical stress granules (Lee *et al.*, 2020). More interestingly, mRNA translation inhibition facilitates formation of PB condensates, whereas increasing the translational activity induces PB disassembly, indicating that the ER contact sites may shuttle mRNAs between the ER and PBs (Lee *et al.*, 2020). In the current study, we observed that the ER/actin network acts as a platform to facilitate trafficking and fusion of the GFP-P granules (Figure 1G; Supplementary Video S1). Given that viroplasms need large amount of BYSMV N to encapsidate newly synthesized genomic/antigenomic RNAs, the BYSMV P-mediated condensates on the ER/actin network opens up the possibility that ER-Viroplasm contact sites are conduits for viral protein and mRNA exchange between the two organelles. Taken together, we propose that the BYSMV P condensates tether on the ER tubules and are dispersed by CK1-mediated phosphorylation. Subsequently, virus transcription is stimulated and viral mRNAs are transported to the ER for efficient translation.

Although the roles of LLPS in viral infections are emerging, multilayered regulatory mechanisms controlling assembly and disassembly of protein condensates remain to be explored. Here, we provide evidence that a plant cytorhabdovirus P protein-mediated LLPS triggers formation of localized viral protein condensates for optimized virus replication. Host CK1-mediated phosphorylation leads to disassembly of viroplasms and hence stimulates virus transcription. Currently, we cannot assemble active viroplasms *in vitro*, because the larger L polymerase protein (234 kDa) is difficult to express and purify from *E. coli*. Therefore, we cannot directly measure the contribution of two P phosphorylation states (P^S5A^ and P^S5D^) to virus replication and transcription *in vitro*. Furthermore, future studies are required to identify the spatiotemporal control mechanisms of host factors including CK1 in viral protein phase separations. Another interesting but unresolved question is how cellular membranes, such as the plasma membrane or ER, regulate the biogenesis and dynamics of viroplasm condensates during virus infection. Nonetheless, these results increases our understanding of the distinct roles of host CK1 and LLPS in uncoupling rhabdovirus transcription and replication.

## Materials and Methods

### Plasmid construction

Based on the pBYSMV-agMR vector (Fang *et al.*, 2019), the inverse PCR product containing an adenine insertion after the start codon of RsGFP was self-ligated to generate the pBY-agMR-RFP vector. To engineer the pGD-GFP-P^WT^, pGD-GFP-P^S5A^ and pGD-GFP-P^S5D^ constructs, the P^WT^, P^S5A^, and P^S5D^ ORFs were amplified from pBYG-P^WT^, pBYG-P^S5A^, and pBYG-P^S5D^ (Gao *et al.*, 2020), respectively, and then introduced into the pGDG vector (Goodin *et al.*, 2002).

The pGD-spGFP1-10-P and pGD-4×spGFP11-P vectors were obtained by replacing the GFP sequence of pGDG-P^WT^ with the cDNA sequences of spGFP1-10 and 4×spGFP11 (Pedelacq *et al.*, 2006; Pedelacq & Cabantous, 2019), respectively. To generate pGD-4×spGFP11-UBC32, the P sequence of pGD-4×spGFP11-P was replaced with the cDNA sequence corresponding to the N-terminal 64 amino acids of UBC32 (AT3G17000) as described previously (Li *et al.*, 2020).

To obtain pET-30a-GFP, pET-30a-GFP-P^WT^, pET-30a-GFP-P^S5A^, pET-30a-GFP-P^S5D^ for recombinant protein expression, the cDNA sequences of 6×His-GFP, 6×His-GFP-P^WT^, 6×His-GFP-P^S5A^ or 6×His-GFP-P^S5D^ were amplified and then inserted into the pET-30a vector for expression of 6×His tagged proteins. The *mCherry* sequence was cloned into pET-32a vector (Novagen) to generate the pET-32a-mCherry vector, into which the BYSMV N ORF was inserted to generate the pET-32a-mCherry-N expression vector.

The pBYSMV-agMR, pGD-N, pGD-P, pGD-L, pGD-VSRs, pGD-ECFP-N, pSuper-mCherry-L, pGDG-NCMV P^WT^, mCherry-HDEL and GFP-ABD2-GFP plasmids have been described previously (Fang *et al.*, 2019). The constructs of pMDC32-NbCK1.3, pGEX-NbCK1.3, pGEX-NbCK1.3^DN^, pGEX-HvCK1.2 and pGEX-HvCK1.2^DN^ have been described previously (Gao *et al.*, 2020), as has pCNX-mRFP (Li *et al.*, 2020).

All sequences were amplified using 2×Phanta Max Master Mix (Vazyme Biotech Co.,Ltd) and inserted into vectors using a ClonExpress Ultra One Step Cloning Kit (Vazyme Biotech Co.,Ltd). Sanger sequencing were performed to confirm sequences. The primers used in this study are listed in Supplementary Table S1.

### Plant materials and protein expression *in vivo*

Four-week-old *N. benthamiana* plants were used for agroinfiltration experiments. The plants were grown in a growth chamber with 16/8 h light/dark cycles and 24°C/20°C (day/night) temperatures. Agroinfiltration experiments for transiently expressing proteins were performed as described previously (Fang *et al.*, 2019). For subcellular localization experiments, *Agrobacterium* harboring plasmids expressing pGD-GFP-P^WT^, pGD-GFP-P^S5A^, pGD-GFP-P^S5D^, pGD-CFP-N, pSuper-L-mCherry, pGD-GFP, pGD-CFP, pSuper-mCherry, pGD-spGFP_1-10_-P (OD_600_, 0.3), pGD-4×spGFP_11_-UBC32 (OD_600_, 0.5), HDEL-mCherry, CNX-mRFP, or pMDC32-NbCK1.3 (OD_600_, 0.1) were mixed with TBSV P19 (OD_600_, 0.1) for infiltration assays. For BYSMV MR assays, *Agrobacterium* harboring pBYS-agMR-RFP, pGD-N, pGD-P/GFP-P^WT^/GFP-P^S5A^/GFP-P^S5D^, pGD-L, pGD-VSRs were diluted to OD_600_ of 0.3, 0.1, 0.1, 0.3, and 0.1, respectively.

### Expression and purification of recombinant proteins

All recombinant plasmids were transformed into *E. coli* BL21 (DE3) cells for recombinant protein expression as described previously (Tong *et al.*, 2021). Briefly, BL21 cells containing different plasmids were grown in 3 mL LB medium with 100 μg/mL Kanamycin overnight at 37°C. The pre-culture was transferred into 1 L LB media with 100 μg/mL Kanamycin and grown to OD_600_ of 0.5 at 37°C, and then induced by 0.5 mM isopropyl β-D-1-thiogalactopyranoside (IPTG) for 18-24 h at 18°C. Bacterial cells were collected and suspended in lysis buffer (30 mM Tris-HCl pH7.5, 500 mM NaCl, 1 mM PMSF and 20 mM imidazole) before being sonicated. After centrifugation (39000 × g, 60 min), the supernatant was flowed throw a column containing 2 mL Ni–nitrilotriacetic acid (NTA) resin equilibrated with lysis buffer. After washing in washing buffer (30 mM Tris-HCl pH7.5, 500 mM NaCl, and 40 mM imidazole), recombinant proteins were eluted with elution buffer (30 mM Tris-HCl pH7.5, 500 mM NaCl, and 400 mM imidazole). For kinase assays in *E. coli*, plasmids encoding 6×His-GFP-P^WT^ with pGEX-NbCK1.3, pGEX-NbCK1.3^DN^, pGEX-HvCK1.2 or pGEX-HvCK1.2^DN^ were co-transformed into *E. coli* Rosetta cells. Expression and purification of 6×His-GFP-P^WT^ were performed as described above (Gao *et al.*, 2020).

### *In vitro* liquid droplet reconstitution assays

All recombinant proteins were centrifuged at 15000 × g for 10 min to remove aggregates. The supernatant concentrations were determined with a NanoDrop spectrophotometer (NanoDrop Technologies) before phase separation assays. All proteins were diluted with buffer (30 mM Tris-HCl pH7.5, 1 mM DTT) to desired protein and salt concentrations. Unless indicated, the final concentration of NaCl was 125 mM and all experiments were performed at room temperature. Phase separation between GFP-P and mCherry-N was conducted by mixing GFP-P with mCherry-N to final concentrations of 10 μM and 6 μM, respectively.

For Cy5-labeled RNAs, 5-10 μg DNA templates of T7 promoter-driven BYSMV trailer sequence served as templates for *in vitro* transcription with the RiboMAX^TM^ Large Scale RNA Production Systems (Promega, P1300) based on the manufacturer’s protocols. Note that the final concentration of ATP, CTP, GTP, UTP, and Cy5-UTP (ApexBio, B8333) in the mixtures were 1.75, 1.75, 1.75, 0.875, and 0.175 mM, respectively. The reactants were mixed gently and incubated at 37°C for 3.5 h, followed by addition of RNase-Free DNase I for 15 min to remove DNA templates. Then, 0.1 volume of 3 M Sodium Acetate (pH 5.2) and 2.5 volumes of 100% ethanol mixtures was added, followed by storage at -20°C for more than 4 h. After centrifugation, the precipitated RNA was washed with 75% ethanol, and suspended in nuclease-free water, and heated at 95°C for 5 min. Phase separation of GFP-P, mCherry-N and Cy5-labeled RNAs was carried out by mixing the indicated proteins and Cy5-labeled RNAs, and diluting with buffer (30 mM Tris-HCl pH7.5, 1 mM DTT) to the desired concentrations.

### Confocal laser scanning microscopy and electron microscopy observations

*N. benthamiana* leaves were agroinfiltrated with *Agrobacterium tumefaciens* containing various constructs and subjected to live-cell imaging at 2-3 dpi with a Leica TCS-SP8 laser scanning confocal microscope. For BYSMV infectivity assays, *N. benthamiana* leaves were observed at 6 days after infiltration with BYS-agMR-RFP. CFP, GFP, mCherry/RFP and Cy5 were excited at 440 nm, 488 nm, 568 nm or 633 nm, and detected at 450-490 nm, 500-540 nm, 585-625 nm or 638-759 nm, respectively. Time-series programs were used to obtain video s. For each video, more than 50 consecutive images were taken at 3- to 5-s intervals and 6 images per second were edited using the Fiji/ImageJ software. Unless indicated, all images were processed using Leica SP8 software. For electron microscopic observations, rBYSMV-RFP-infected barly stems were fixed and embedded in Spurr’s resin, and ultra-thin sections were observed with a Hitachi TEM system.

### Fluorescence recovery after photobleaching (FRAP)

FRAP were performed with a Leica SP8 laser scanning confocal microscope (63×/100× oil objective, PMT detector). *N. benthamiana* leaves were agroinfiltrated to express the GFP-P^WT^ protein, and then subjected to living-cell imaging at 2-3 dpi. Note that *N. benthamiana* leaves were treated with 10 mM LatB (Abcam) to inhibit trafficking of GFP-P^WT^ granules at 3 h before FRAP assays. Then the GFP-P granules were bleached three times with a 488-nm laser at 100% laser power and time-lapse modes were used to collect recovery images. For *in vitro* FRAP assays, droplets were bleached once with a 488-nm laser at 50% laser power with ≥ 6 samples. The Fiji/ImageJ software was used to measure fluorescence intensity and the recovery curves were carried out with Graphpad Prism8 software.

### Western blotting analysis

Total proteins were isolated from *N. benthamiana* leaf tissues in extraction buffer (100 mM Tris-HCl, pH 6.8, 20% glycerol, 4% SDS, 0.2% bromphenol blue, 5% β-mercaptoethanol). Total proteins were separated in a 4–15% SDS–PAGE gradient and transferred to nitrocellulose membrane (GE Healthcare Life Science). Membranes were blocked with 5% (m/v) skimmed milk powder at room temperature for 1 h and then incubated with primary antibodies at 37°C for 1 h. After washing three times, membranes were incubated with secondary antibodies at 37°C for 1 h. Antibodies against RFP (1:3000), P (1:3000), GST (1:5000), GFP (1:5000; MBL, 598), Flag (1:5, 000; Sigma, F1804) were used for protein detection. Goat anti-rabbit IgG (EASYBIO, BE0101) and goat anti-mouse IgG horseradish peroxidase conjugate (Bio-Rad, 170-6516) were used as secondary antibodies. After addition of NcmECL Ultra stabilized peroxide reagent (NCM Biotech, P10300B), chemiluminescence of membranes was detected with a Biomolecular Imager (Azure biosystems, Inc.).

### BYSMV P granule sedimentation assay

Granule purification assays were performed as described previously (Hubstenberger *et al.*, 2017). *N. benthamiana* leaves (0.3 g) expressing GFP-P^WT^, GFP-P^S5A^ and GFP-P^S5D^ were ground in liquid nitrogen and suspended in 600 μL lysis buffer (50 mM Tris-HCl, pH 7.5, 1 mM EDTA, 150 mM NaCl, 1 mM DTT, 0.2% Triton X-100). After centrifuging at 13000 × g for 10 min, the supernatants were added 600 μL extraction buffer (100 mM Tris-HCl, pH 6.8, 20% glycerol, 4% SDS, 0.2% bromphenol blue, 5% β-Mercaptoethanol) and used as soluble protein samples. The pellets were suspended in 1 mL lysis buffer and centrifuged at 6000 × g for 10 min to deplete free GFP-P^WT^, GFP-P^S5A^ and GFP-P^S5D^, and the pellets were resuspended in 200 μL extraction buffer for use as pellet samples. For input samples, 0.1 g of grounded *N. benthamiana* leaves were suspended in 500 μL extraction buffer. All the input, supernatant, and pellet samples were used to detect GFP-P accumulation by western blotting analyses with antibodies against BYSMV P.

### 1, 6-hexanediol (HEX) treatment assay

The 1, 6-hexanediol reagent (Sigma-Merck, 240117) were heated to 45°C and diluted to 50% in water as stock solution. For *in vivo* assays, 50% 1, 6-hexanediol was diluted to 2.5%, 5%, and 10% in water, and infiltrated into *N. benthamiana* leaves expressing GFP-P^WT^ at 2 dpi. After 5 min, *N. benthamiana* leaves were observed under a Leica SP8 laser scanning microscope. For *in vitro* assays, the GFP-P^WT^ droplets were treated with 5% 1, 6-hexanediol for 1 minute and then were observed under a scanning microscope.

In the HEX-treated SBPHs assays, rBYSMV-GFP infected barley leaves were homogenized in buffer (1 mM MnCl2, 10 mM Mg (CH3COO)2, 40 mM Na2SO3 and 100 mM Tris-HCl, pH 8.0) and centrifuged at 12,000 g for 20 min at 4°C. The suspension was divided into two equal parts, which were added 3% 1, 6-hexanediol and ddH2O, respectively. Then, 18.6 nL of the mixture was injected into the second-instar nymph using a Nanoinject II auto-nanoliter injector (Drummond Scientific). The injected nymphs were reared on fresh rice seedlings and photographed with an Olympus stereomicroscope SEX16 (Olympus, Tokyo, Japan) at 3- and 5- dpi.

In the HEX-treated barley assays, rBYSMV-GFP-infected SBPHs were transferred onto 4-day-old healthy barley leaves for a 10-h inoculation period. Then, the infected barley leaves were infiltrated with 3% HEX or buffer. More than five barley plants were used in each treatment. At 10 dpi, the Biomolecular Imager (Azure biosystems, Inc.) were used to detect GFP fluorescence of rBYSMV-GFP-infected plants. The inoculated whole barley plants were used for detection of viral accumulation by Western blotting assays with specific antibodies against BYSMV N or GFP.

### *In vitro* dephosphorylation assays

The phosphorylated GFP-P^WT^ protein, purified from *E. coli* Rosetta cell containing pGEX-NbCK1.3 or pGEX-HvCK1.2 plasmid, was incubated with 10 U/ul Lambda Protein Phosphatase(New England Biolabs, #P0753) at 30°C for 30 minutes. Samples were resuspended in extraction buffer (100 mM Tris-HCl, pH 6.8, 20% glycerol, 4% SDS, 0.2% bromphenol blue, 5% β-mercaptoethanol) and analyzed by SDS-PAGE.

### Quantitative reverse transcription PCR (RT-qPCR) assays

The RT-qPCR assay was performed as described previously (Gao *et al.*, 2020). Briefly, total RNA isolated from plants was firstly treated with DNase I (Takara) to remove DNA contamination. Next, 2.5 μg total RNA was used as a template for reverse transcription by HiScript II Reverse Transcriptase (Vazyme Biotech Co., Ltd) with primers oligo (dT)/BYS-RT-F and qNbEF1α-R. Quantitative PCR (qPCR) was performed using SsoFast EvaGreen Supermix (Bio-Rad) on CFX96 Real-Time system (Bio-Rad). The *EFIA* gene was used as an endogenous control. Three independent biological replicates were used for biological statistics analysis. All the primers used in this study are provided in Supplementary Table 1.

### Prediction of IDRs

The IDRs were predicted with the online tool PONDR (http://www.pondr.com/) with default parameters.

### Quantification and statistical analyses

Images were analyzed with Fiji/ImageJ software. At least three independent replicates were used for all experiments, and statistical analyses were done using the Graphpad Prism8 software. Statistical significance was assessed by unpaired two-tailed Student’s t test.

## Acknowledgments

We thank Prof. Jialin Yu, Chenggui Han, and Yongliang Zhang for their helpful discussion, and Yan Zhang for her technique assistance. We thank Prof. Caiji Gao (South China Normal University) for providing the CNX-mRFP and spGFP1-10- UBC32 plasmids. We thank Prof. Xiaorong Tao (Nanjing Agricultural University) for his gift of the GFP-ABD2-GFP and HDEL-mCherry plasmids. This work was supported by the Natural Science Foundation of China 31872920 and 31571978 (XBW) Chinese Universities Scientific Fund 2021TC063 (XBW).

## Author contributions

X.F. performed the majority of the experiments, assisted by Q.G., Y.Z., J.Q., D. G. and W.X. X.F. and X.W. analyzed the data. X.W and X.F. wrote the manuscript, assisted by Y.W. and D.L. All authors discussed the results and commented on the manuscript.

---

**Figure supplement 1.**
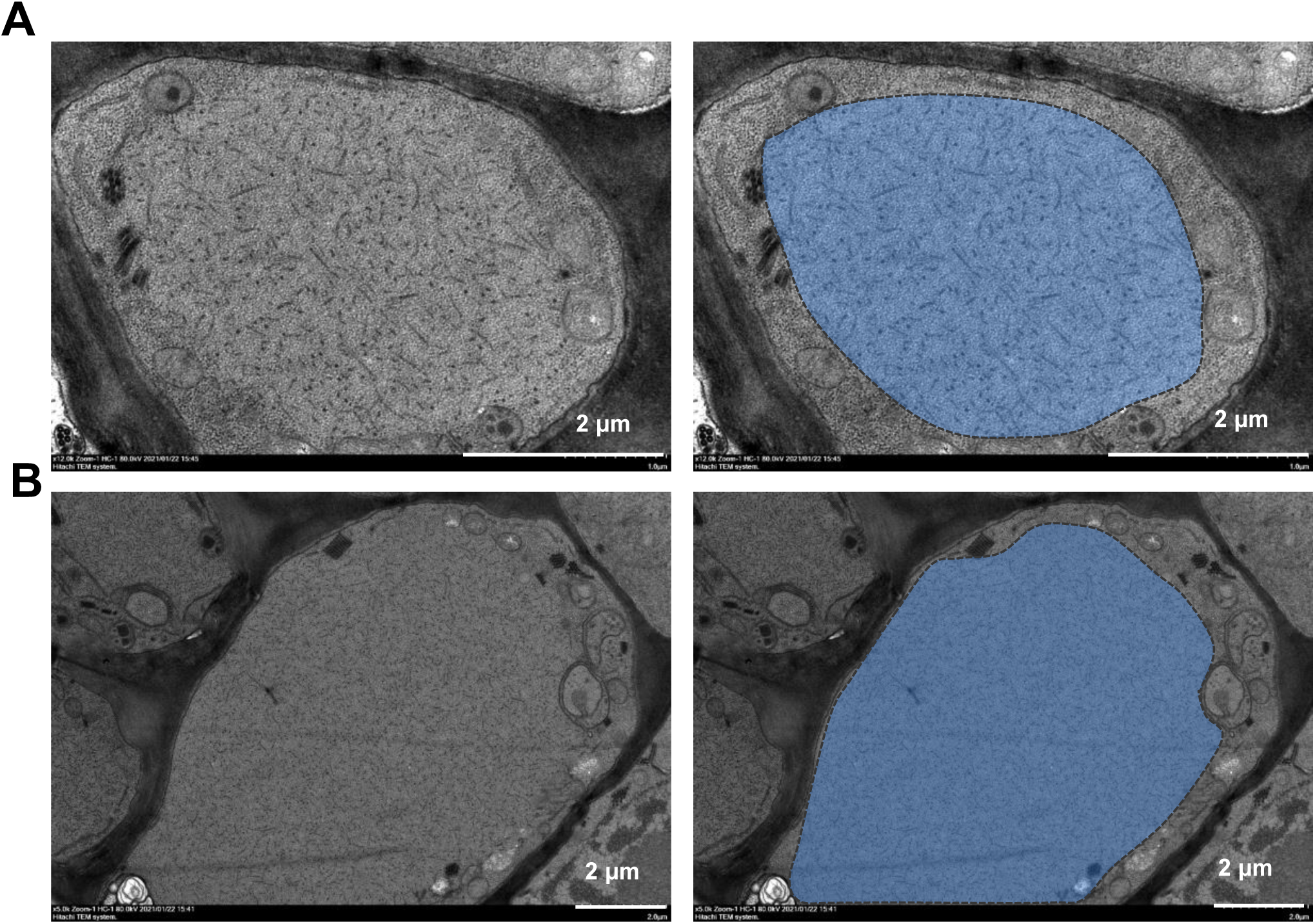
EM characterization of BYSMV viroplasms in the ultrastructural aspects of barley leaves infected by BYSMV. **(A)** and **(B)**, transmission electron microscopy characterization of the BYSMV viroplasm in barley leaves infected by BYSMV at 25 dpi. The electron dense granular structure of the BYSMV viroplasm was highlighted by blue color in the right panel. Scale bars, 2 μm.

**Figure supplement 2.**
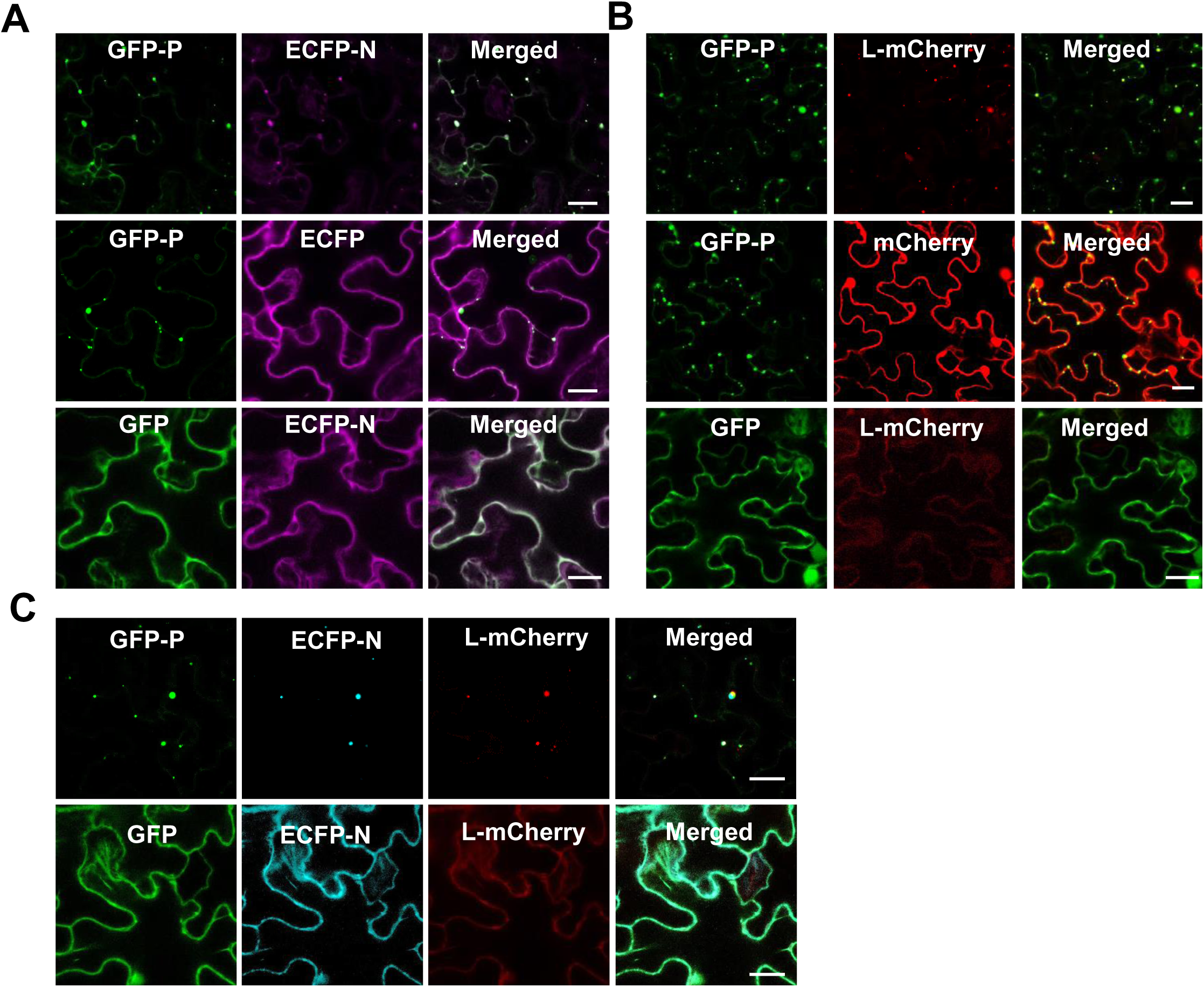
BYSMV P forms spherule granule recruiting N and L proteins. **(A)** Confocal images showing subcellular localization of GFP-P/GFP and ECFP-N/ECFP transiently expressed in different combinations in epidermal cells of *N. benthamiana* leaves. Scale bars, 20 μm. **(B)** Confocal images showing subcellular localization of GFP-P/GFP and L-mCherry/mCherry transiently expressed in different combinations in epidermal cells of *N. benthamiana* leaves. Scale bars, 20 μm. **(C)** Confocal images showing subcellular localization of CFP-N and L-mCherry co-expressed with GFP or GFP-P in epidermal cells of *N. benthamiana* leaves at 2 dpi. Scale bars, 20 μm.

**Figure supplement 3.**
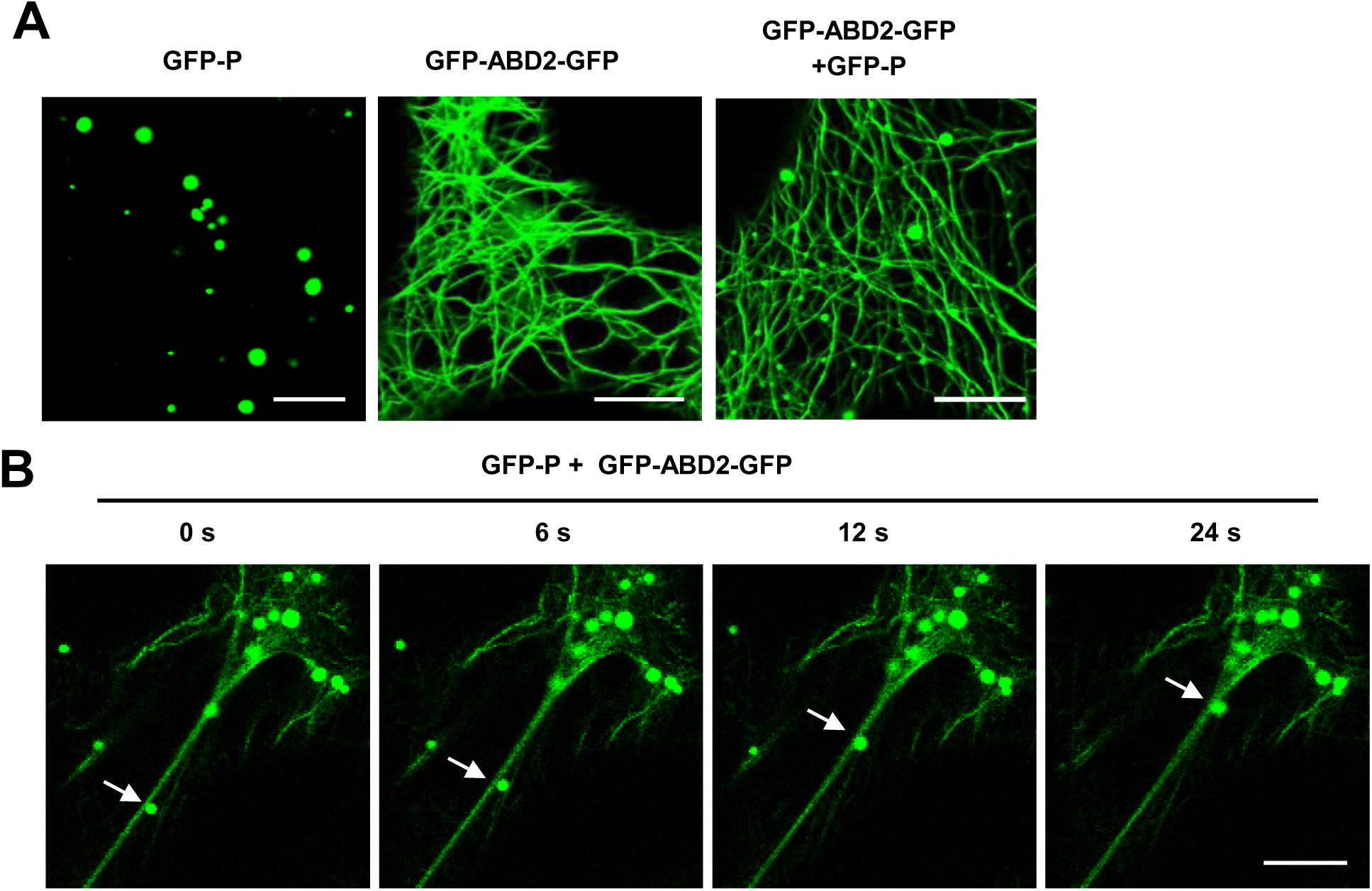
GFP-P moved along the actin filaments. **(A)** Subcellular localization of GFP-P or GFP-ABD2-GFP expressed alone or together in *N. benthamiana* leaf epidermal cells at 2 dpi. Scale bars, 10 μm. (B) Subcellular localization of GFP-P granules with GFP-ABD2-GFP-labeled actin filaments at 2 dpi. White arrows indicate that GFP-P granules moved along the actin filaments. Scale bar, 10 μm.

**Figure supplement 4.**
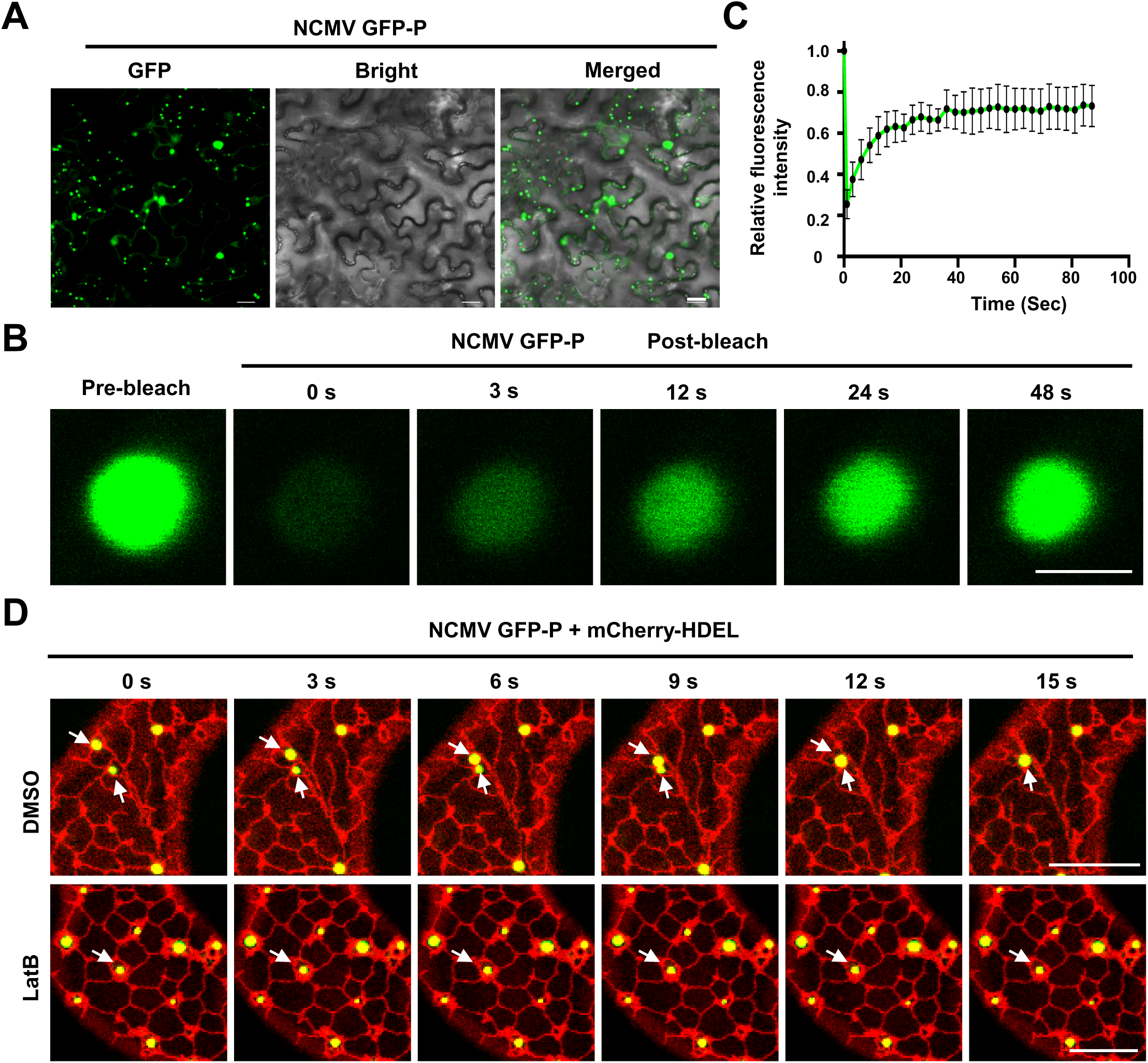
NCMV P forms liquid structures *in vivo*. **(A)** Confocal images showing subcellular distribution of NCMV GFP-P transiently expressed in *N. benthamiana* leaf epidermal cells at 2 dpi. Scale bar = 20 μm. **(B)** Representative FRAP images of NCMV GFP-P granules transiently expressed in epidermal cells of *N. benthamiana* leaves at 2 dpi. Leaves were treated with 10 μM LatB to inhibit traffic of NCMV GFP-P granules at 3 h before FRAP imaging. Scale bars, 2 μm. **(C)** Representative FRAP recovery curves of NCMV GFP-P granules as shown in panel B. The intensity of each body was normalized against their pre-bleach fluorescence. Data were expressed as the mean ± SD of 10 foci. **(D)** Time-lapse confocal micrographs showing the localization of NCMV GFP-P and mCherry-HDEL transiently expressed in *N. benthamiana* leaf epidermal cells at 2 dpi. Leaves were treated with DMSO or 10 μM LatB at 3 h before imaging. White arrows indicate that GFP-P granules undergo fusion. Scale bars, 10 μm.

**Figure supplement 5.**
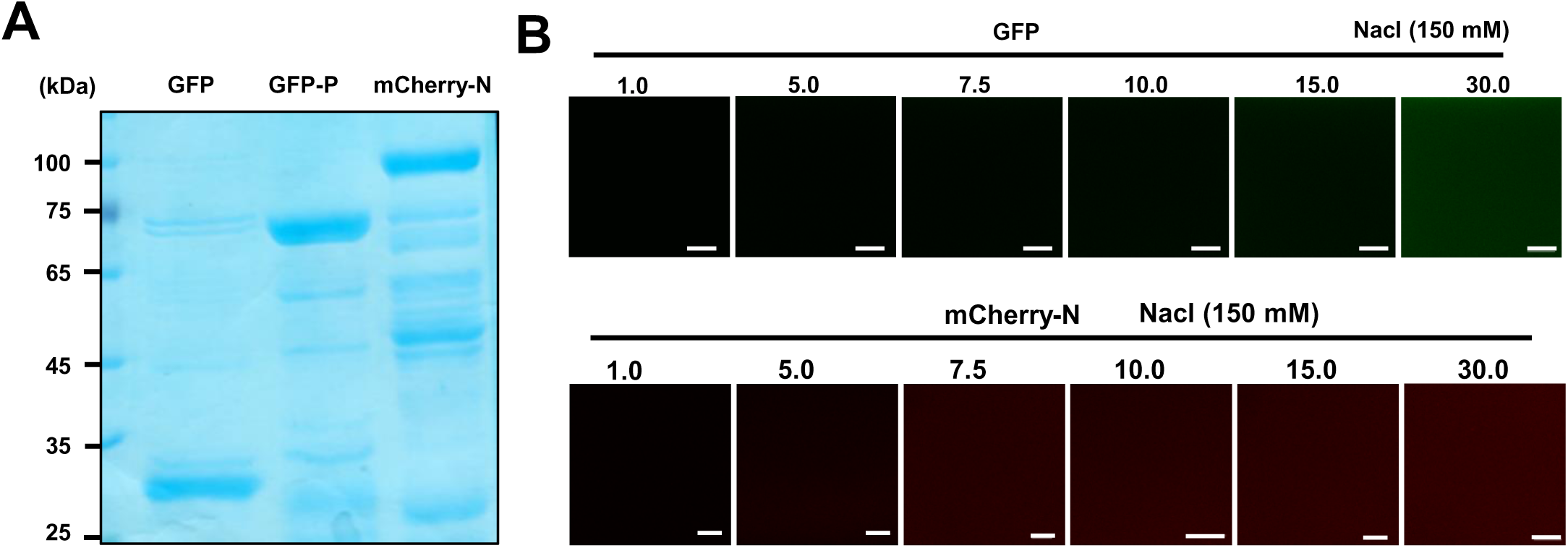
GFP and BYSMV mCherry-N are deficient in forming liquid droplets *in vitro*. **(A)** SDS-PAGE showing purified GFP, GFP-P, and mCherry-N proteins. **(B)** Confocal images showing that neither GFP nor mCherry-N formed droplets at the concentration of 1-30 μM in 150 mM NaCl. Scale bars, 20 μm.

**Figure supplement 6.**
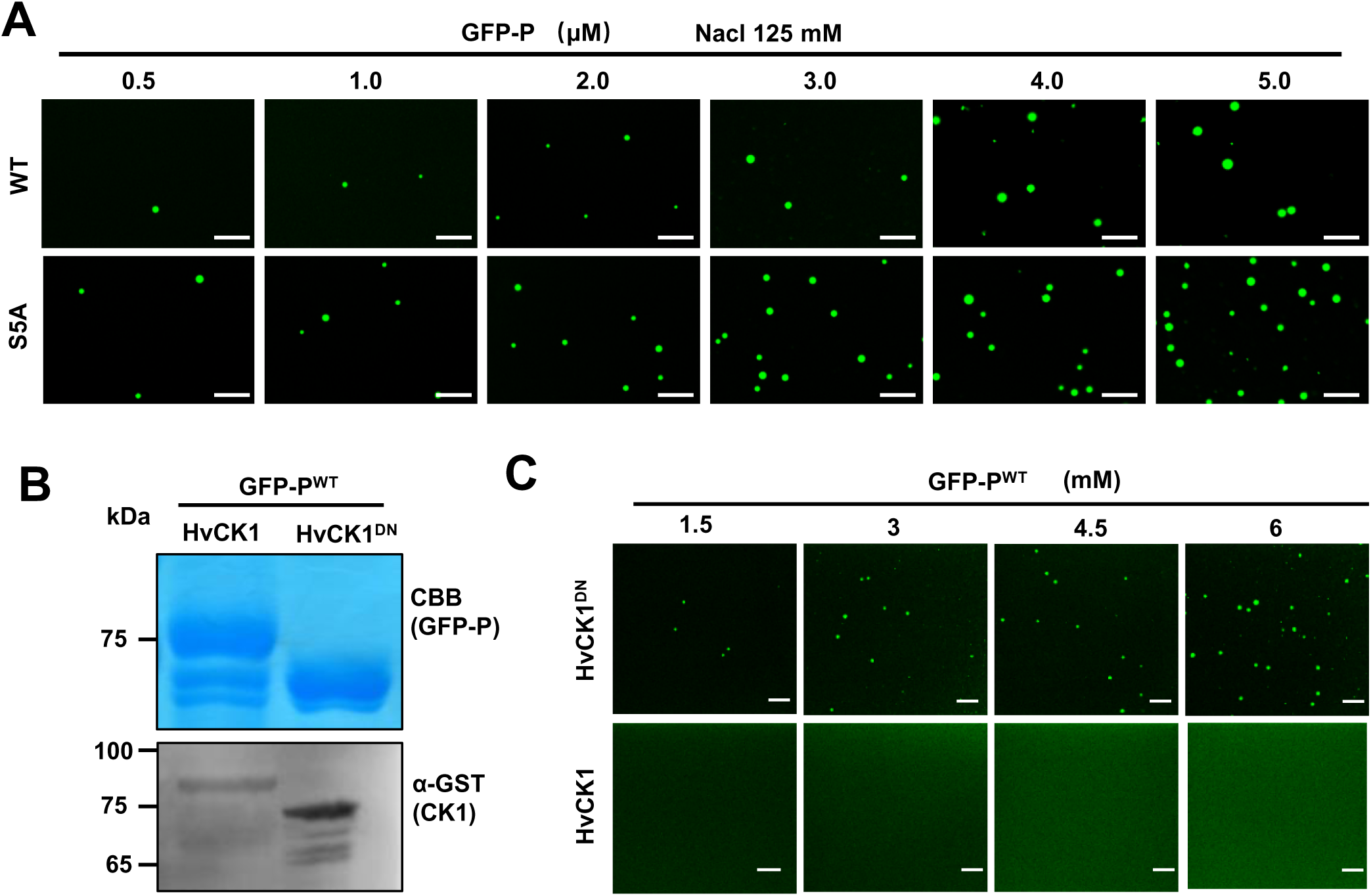
Phase separation of BYSMV P is inhibited by P protein phosphorylation. **(A)** Confocal images showing that GFP-P^WT^ and GFP-P^S5A^ formed droplets at different concentrations and 125 mM NaCl. Scale bars, 20 μm. **(B)** SDS-PAGE showing purified GFP-P^WT^ purified from *E. coli* co-expressing HvCK1 or HvCK1^DN^. Accumulation of GST-tagged HvCK1 or HvCK1^DN^ was examined by Western blotting analyses with anti-GST antibodies. **(C)** Confocal images showing droplet formation of GFP-P^WT^ purified from *E. coli* co-expressing HvCK1^DN^. GFP-P^WT^ was diluted to different concentration in 125 mM NaCl. Scale bars, 20 μm.

**Figure supplement 7.**
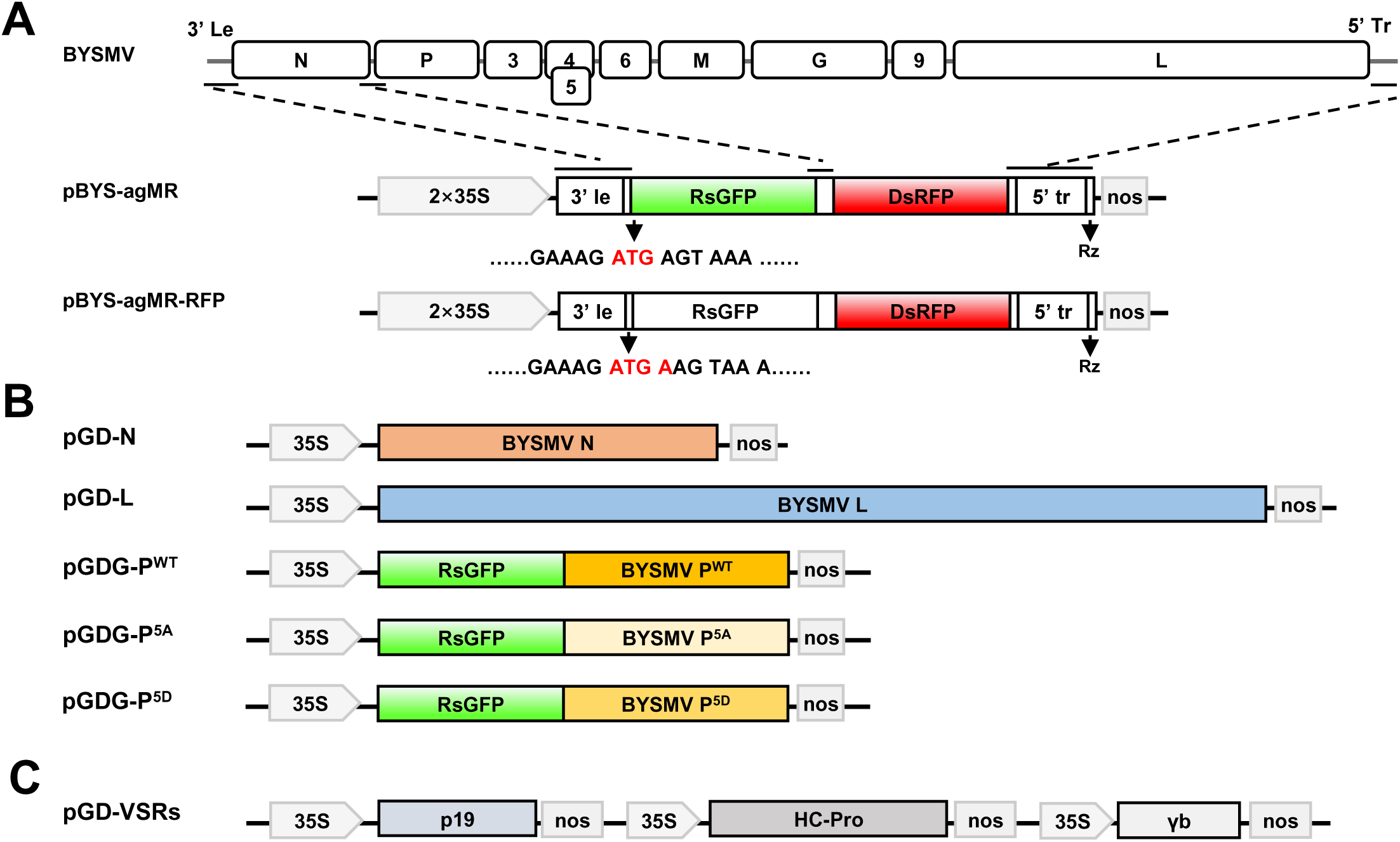
Illustration of the binary vectors used to rescue BYS-agMR-RFP infection in epidermal cells of *N. benthamiana* leaves. **(A)** In the pBYSMV-agMR plasmid, the ORFs of BYSMV N and P were replaced with GFP and RFP, respectively, and were flanked by 3′ leader (le) and 5′ trailer (tr) sequences outside the N gene and the L gene. Then, the reporter cassette of agMR was inserted between the truncated CaMV double 35S promoter and the hepatitis delta (HDV) ribozyme sequence (RZ). Based on the pBYS-agMR vector, the pBYS-agMR-RFP vector was generated by inserting a nucleotide A after the start codon of GFP to silence expression of GFP. **(B)** The ORFs of N, L, and GFP-P^WT^/GFP- P^S5A^/GFP-P^S5D^ were inserted into the pGD vector for expression of N, L, and GFP-P^WT^/GFP- P^S5A^/GFP-P^S5D^. **(C)** Three virus suppressors of RNA silencing (VSRs), including the Tomato bushy stunt virus p19, the Tobacco etch virus HC-Pro, and the Barley stripe mosaic virus γb, were individually inserted into the pGD vector, and then these cassettes were combined into one plasmid for expression of three VSRs simultaneously. Note that BYS-agMR-RFP was rescued by co-expression of pBYS-agMR-RFP, N, L, P derivative, and VSRs in *N. benthamiana* leaves.

**Supplementary table 1.**
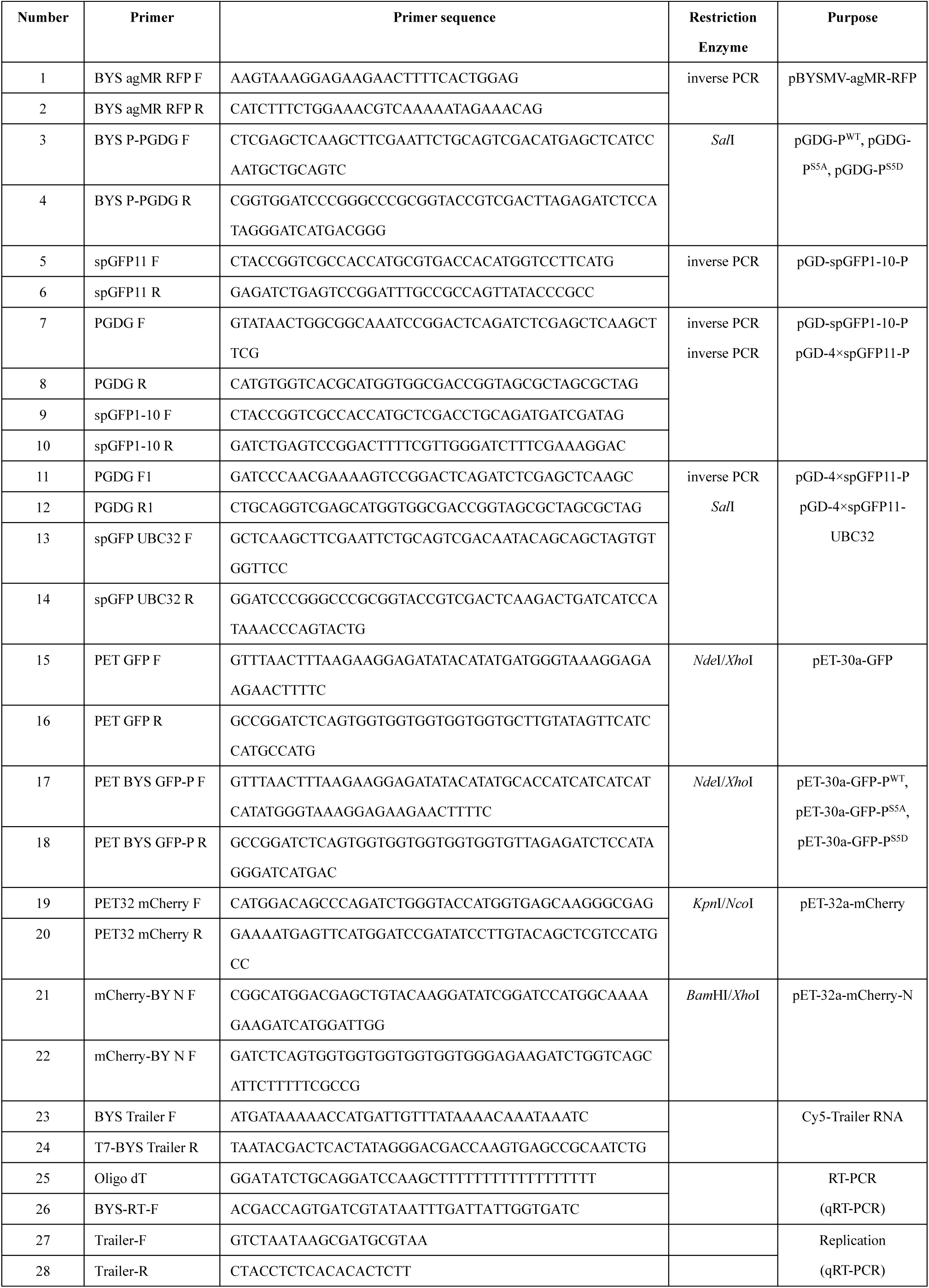
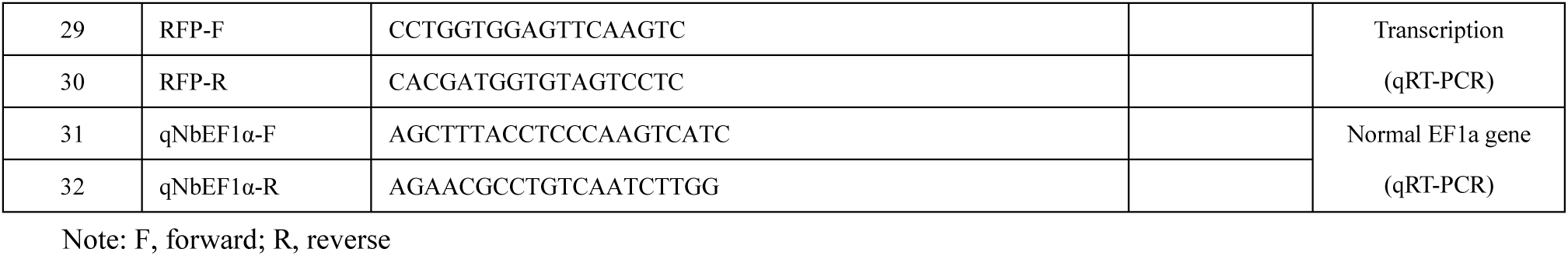
Primers used in this study

## Supporting Information

**Supplementary Video S1.** Trafficking and fusion video of BYSMV GFP-P graunles in epidermal cells of *N. benthamiana* leaves at 2 dpi.

**Supplementary Video S2.** Representative FRAP video of BYSMV GFP-P granules transiently expressed in epidermal cells of *N. benthamiana* leaves at 2 dpi.

**Supplementary Video S3.** Time-lapse confocal video showing the localization of GFP-P and mCherry-HDEL expressed in *N. benthamiana* leaf epidermal cells at 2 dpi.

**Supplementary Video S4.** Time-lapse confocal video showing the localization of GFP-P and mCherry-HDEL expressed in *N. benthamiana* leaf epidermal cells at 2 dpi.

**Supplementary Video S5.** Trafficking and fusion video of NCMV GFP-P graunles in epidermal cells of *N. benthamiana* leaves at 2 dpi.

**Supplementary Video S6.** Representative FRAP video of NCMV GFP-P granules transiently expressed in epidermal cells of *N. benthamiana* leaves at 2 dpi.

**Supplementary Video S7.** Representative FRAP video of BYSMV GFP-P droplets *in vitro*.

**Supplementary Video S8.** Representative video showing fusion of BYSMV GFP-P droplets *in vitro*.

